# Periodic shifts in viral load increase risk of spillover from bats

**DOI:** 10.1101/2023.09.06.556454

**Authors:** Tamika J. Lunn, Benny Borremans, Devin N. Jones, Maureen K. Kessler, Adrienne S. Dale, Kwe C. Yinda, Manuel Ruiz-Aravena, Caylee A Falvo, Dan Crowley, James O. Lloyd-Smith, Vincent J. Munster, Peggy Eby, Hamish McCallum, Peter Hudson, Olivier Restif, Liam P. McGuire, Ina L. Smith, Bat One Health Group, Raina K. Plowright, Alison J. Peel

## Abstract

Prediction and management of zoonotic pathogen spillover requires an understanding of infection dynamics within reservoir host populations. Transmission risk is often assessed using prevalence of infected hosts, with infection status based on the presence of genomic material. However, detection of viral genomic material alone does not necessarily indicate the presence of infectious virus, which could decouple prevalence from transmission risk. We undertook a multi-faceted investigation of Hendra virus shedding in *Pteropus* bats, combining insights from virus isolation, viral load proxies, viral prevalence, and longitudinal patterns of shedding, from 6,151 samples. In addition to seasonal and interannual fluctuation in prevalence, we found evidence for periodic shifts in the distribution of viral loads. The proportion of bats shedding high viral loads was higher during peak prevalence periods during which spillover events were observed, and lower during non-peak periods when there were no spillovers. We suggest that prolonged periods of low viral load and low prevalence reflect prolonged shedding of non-infectious RNA, or viral loads that are insufficient or unlikely to overcome dose barriers to spillover infection. These findings show that incorporating viral load (or proxies of viral load) into longitudinal studies of virus excretion will better inform predictions of spillover risk than prevalence alone.

**Significance statement:** We present a comprehensive analysis of a high-profile bat-virus system (Hendra virus in Australian flying-foxes) to demonstrate that both prevalence and viral loads can shift systematically over time, resulting in concentrated periods of increased spillover risk when prevalence and viral loads are high. We further suggest that prolonged periods of low-prevalence, low-load shedding may not reflect excretion of infectious virus, resolving the outstanding puzzle of why spillovers have not been observed during periods of low off-season prevalence in subtropical Australia, or more frequently in tropical Australia despite consistent low-prevalence shedding. The consideration of viral loads (or proxies of viral load) along with prevalence may improve risk inference from longitudinal surveys of zoonotic viruses across wildlife reservoir hosts.

## Introduction

Emergence of zoonotic infections with epidemic potential is a public health concern (Lloyd-Smith et al. 2009). A major challenge for the design and implementation of pre-emptive strategies emerges from the complex, multilayered nature of cross-species transmission (Plowright et al. 2017). The pathogen, its reservoir host(s), environment, and potential novel hosts are connected dynamically in space and time, with intermittent opportunities for cross-species transmission when conditions align. The dynamics of pathogen circulation in reservoir host populations are an important determinant of spillover risk, and are usually assessed through the prevalence of infection in space and time (Plowright et al. 2019). Although viral load has been highlighted as a potentially crucial factor, it has rarely been considered in empirical studies of spillover risk (Lunn et al. 2019).

For pathogens where prolonged viral shedding can be detected after the disappearance of active infection, the estimation of prevalence based on detection or non-detection of genomic material can decouple presence of pathogen from the risk of transmission (Griffin 2022). For example, RNA shedding weeks to months after the disappearance of infectious virus has been observed for measles virus, severe acute respiratory syndrome coronaviruses (SARS-CoV and SARS-CoV-2), and Ebola virus (Chan et al. 2004; Lin et al. 2012; Sissoko et al. 2017; Bullard et al. 2020). Biologically, prolonged RNA shedding can occur because the immune response utilizes a multitude of mechanisms to neutralize viruses and prevent replication, without necessarily eliminating degraded nucleic/ribonucleic acid (Atkinson and Petersen 2020, reviewed in Griffin 2022). Non-infectious genomic material can therefore be shed as non-enveloped, relatively stable, nucleoprotein complexes in the absence of active infection and production of infectious virions (Compton et al. 2004; Griffin 2022). Practically, assays target specific segments of RNA, so even with extensive RNA degradation, there may be limited impact on the assay’s ability to detect RNA. This creates challenges for assessment and prediction of infectious virus excretion, or virus doses that might result in spillover.

Viral load in humans has been used to estimate transmissibility (e.g., HIV; Tovanabutra et al. 2002), and to predict patient outcome (e.g., Ebola Virus Disease; Matson et al. 2022). Recent research prompted by the COVID-19 pandemic has further demonstrated the utility of viral load (or proxies of viral load) in predicting trajectories of epidemics in epidemiological modelling (e.g., Hay et al. 2021; Funk and Abbott 2022). For wildlife studies, integration of viral load with prevalence will provide a better representation of how infectious virus is distributed across a landscape, and the dose available for recipient hosts at a given point in space and time (Plowright et al. 2017; Becker et al. 2023), but to-date has been overlooked in viral surveillance and forecasting. Quantitative measurement of viral shedding intensity – particularly through the calculation of genome copy numbers as a proxy of viral load – is a valuable but underexplored addition to population-level models informing predictions of spillover risk from wildlife.

Hendra virus is a bat-borne paramyxovirus (genus: *Henipavirus*) that causes severe disease and death in horses and humans in eastern Australia (Field et al. 2001; Playford et al. 2010). Equine cases are reported annually with detection of clusters in some years (Eby et al. 2023). Sixty-seven spillover events, involving 109 confirmed or suspected cases in horses, and 7 confirmed human cases, have been reported since the virus was first detected in 1994, including two detections of a recently discovered variant (HeV-g2) in horses (Queensland Government 2023; Annand et al. 2022). As a result of high virulence and recurrent spillover of henipaviruses, the World Health Organization has listed henipaviral diseases (including Hendra virus disease in Australia and Nipah virus disease in Asia) as one of the highest priority diseases for research (World Health Organization 2022).

Despite the alignment of bats and horses across most of eastern Australia, spillovers have only been detected in coastal New South Wales and Queensland. Most occur in winter in the subtropics and in clusters that occur with inter-annual variation (Plowright et al. 2015; Eby et al. 2023). In part, this may be explained by increased prevalence of viral shedding from bats (measured as the proportion of positive PCR detections from pooled samples of urine) in winter, and the varying magnitude and timing of shedding peaks (Field et al. 2011; Field et al. 2015; Páez et al. 2017; Becker et al. 2023). Previous work also suggests periods of low off-season prevalence in roosts from subtropical Australia, and consistent low-prevalence shedding from roosts in tropical Australia (Field et al. 2015; Páez et al. 2017). Why these prolonged periods of shedding do not translate to spillover has been a persistent puzzle. Whether cumulative, low-level shedding over a long time presents a similar risk as high-level shedding over a short time also remains unknown (Páez et al. 2017). Incorporation of viral load (or viral load proxies) into longitudinal studies of virus excretion may better inform prediction of periods and places of higher spillover risk.

To refine predictions and manage risk, we sought to test whether temporal variations in viral loads are more predictive of spillover risk than measures of prevalence alone. For this, we sampled five flying-fox roost sites in subtropical Australia for three and a half years, and screened for excretion of Hendra virus (HeV-g1). We incorporated information on viral load proxies (Ct and genome copy numbers) alongside virus isolation data to estimate the amount of infectious virus shed from flying-fox populations. We focused on roosts with ecological characteristics linked to high risk of spillover (Becker et al. 2023; Eby et al. 2023), and used an optimized pooled sampling method (Giles et al. 2018) to improve accuracy of prevalence estimates. Our new dataset and multi-faceted analyses provide insights into observed but unexplained patterns of shedding and spillover, and will contribute to the future management of viral spillover risk.

## Methods

We conducted our study from July 2017 through September 2020 at five roosts spanning south-east Queensland to north-east New South Wales (“Redcliffe”, “Sunnybank”, “Toowoomba, “Burleigh”, and “Clunes”; Figure 1). These roosts were selected because they have attributes associated with higher risk of spillover: 1) continuous occupation by black flying-foxes (*Pteropus alecto*); 2) recently established as overwintering; and 3) highly restricted access to native winter food sources (Becker et al. 2023; Eby et al. 2023). The commencement of sampling was staggered across sites, with all sites incorporated by April 2018. Sampling of roosts occurred approximately monthly, with most sampled within the same week each month. For analyses performed on contemporaneous monthly sessions across aggregated sites, clusters of sessions were defined such that all sessions within the cluster were within 14 days of each other. Five Hendra virus spillover events occurred in the study area during our sampling period, with two falling within the foraging radius of our study sites (a single study site, “Clunes”) (Eby et al. 2023).

**Figure 1:**
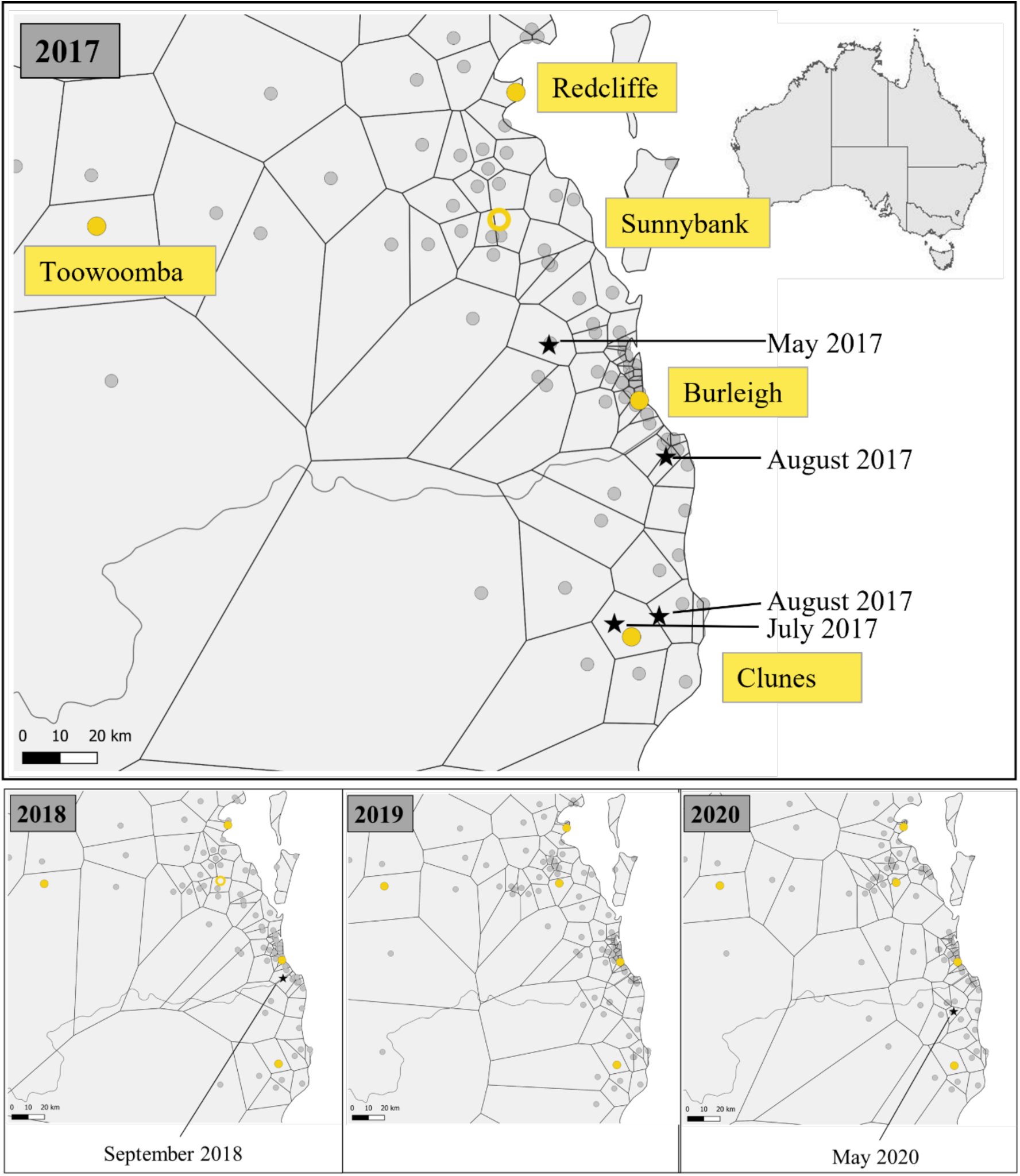
Location of winter-occupied flying-fox roosts (grey circles) and Hendra virus spillover events (black stars) in the study area, per study year (2017-2020). Tessellation boundaries show the nearest winter-occupied flying-fox roost for any given location. Roosts sampled for the present study are shown in yellow. Note that the roost site “Sunnybank” was not occupied over winter in 2017. The location of the roost is shown as a hollow circle on the 2017 and 2018 panels for reference. Flying-fox roost locations obtained from Eby et al. (2023).

### Sample collection

At each site, we collected pooled urine samples from beneath roosting trees using plastic sheets (0.9 x 1.3 meters) distributed before sunrise (∼4AM to 6AM Australian Eastern Standard Time), following an optimized sampling approach (Appendix S3). Sample collection began once bats had returned to the roost and concluded within 6 hours. We pooled urine on each sheet into a single sample (roughly 20 urine droplets to a volume of ∼2ml), and aliquoted up to three cryovials for Hendra virus testing containing: AVL buffer (Qiagen) (target 140 µl of urine into 560 µl buffer), viral transport medium (VTM; Munster et al. 2009) (between 200-1000 µl of urine into 1000 µl of VTM), and no buffer. Samples were transported in a CryoShipper (< -80°C) and stored at -80°C. An average of 40 pooled samples were collected per session. Note that sheet placement was targeted for black flying-foxes (*P. alecto*), but grey-headed (*P. poliocephalus*) and, rarely, little red flying foxes (*P. scapulatus*) were also sampled. Urine was not collected from sheets if the urine had completely evaporated or was contaminated with feces. Rain-affected sampling events were discarded and re-scheduled.

### qRT-PCR screening of Hendra virus

We used the QIAamp Viral RNA Kit and QIAcube HT automated system (Qiagen) to extract RNA. RNA was eluted in 150μl of TE buffer. Hendra virus genotype-1 (HeV-g1) duplex qRT-PCR assay was used for the detection of viral RNA (Appendix S3). In the qRT-PCR, 10μl RNA, 900nM and 250nM final concentrations of primer and probe (Appendix S3), respectively, were tested with 10μl of TaqMan Fast Virus 1-Step master mix on QuantStudio 6 flex Real-Time PCR instrument (Applied Biosystems).

Thermocycling conditions are given in Appendix S3. Positive standards (with known genome copy numbers previously quantified using droplet digital PCR) were run in parallel with samples at ten-fold dilutions to construct a standard curve and calculate genome copy numbers/mL of each positive sample. We considered genome copies and Ct values to be proxies for viral load, and use the term “viral load” throughout.

Prior to statistical analyses we assessed effects of methodology and environment on virus detection, including sample volume, collection time, temperature at collection, and whether sample had begun to evaporate or was in direct sun at collection (Appendix S4). We did not observe systematic effects of these covariates, consistent with prior expectations (Munster et al. 2009). However, sample buffer (AVL, VTM, or none) did impact the likelihood of virus detection (Appendix S4).

### Virus isolation

Samples are traditionally deemed positive for Hendra virus with detection at Ct≤40 in qRT-PCR assays, but detection of genetic material does not discriminate infectious from non-infectious RNA (Jaafar et al. 2021). We compared isolation success (indicative of functionally infectious virus) with Ct from qPCR positive samples. Virus isolation was performed on samples obtained in virus transport media (VTM). Vero E6 cells in a 24-well plate were inoculated with 250 µl original undiluted sample and a 1:10 dilution thereof in different wells. Diluted and undiluted samples were plated in duplicates. Plates were centrifuged for 30 minutes at 1000 rpm and incubated for 30 minutes at 37°C and 5% CO_2_. The inoculum was removed and replaced with 500 µl DMEM containing 2% FBS, 50 U/ml penicillin and 50 μg/ml streptomycin. Two and three days after inoculation, cytopathic effect (CPE) was scored. Blind passage was performed on samples without CPE as above; other than that, plates were not spun. Supernatants from plates with CPE present were analyzed via qPCR for Hendra virus RNA to rule out other causes of CPE. The obtained virus culture was categorized into binary outcomes (CPE positive or negative) and plotted against Ct. Spearman correlation was used to determine association between Ct and CPE. These primary data were combined with Hendra virus isolation data from published horse, human and flying fox studies (Playford et al. 2010; Barr et al. 2015), to facilitate investigation into the relationship between Ct value and viral isolation.

### Prevalence and viral load distribution

We assessed whether viral loads are randomly distributed over time, or whether there are periods during which proportionally more individuals shed high viral loads, using a permutation analysis. First, we estimated a proxy for prevalence as the proportion of pooled urine samples positive for Hendra virus, with temporal dynamics estimated from a generalized additive model (GAM) fitted to time. Models used a binomial distribution with a logit-link, with prevalence modeled as a natural cubic spline with 8 degrees of freedom. Model fits were repeated for different Ct thresholds, ranging from Ct 28 (∼100,000 genome copies/ml) to Ct 39 (∼100 genome copies/ml). To allow comparison between Ct threshold values, prevalence was normalized by dividing by maximum prevalence across the dataset for each threshold. Normalized prevalence should be interpreted as a relative value, where high values indicate when prevalence is high relative to other values in the time series for each Ct threshold. For non-normalized prevalence values at each Ct threshold, see Appendix S1.

Permutations randomly shuffled Ct values of positive samples to re-estimate prevalence dynamics with randomized time, with Ct values shuffled prior to the application of Ct thresholds. Permuted prevalence dynamics estimated using the conventional Ct threshold (Ct 40, ∼60 genome copies/ml) are thus the same as for the observed data. Prevalence dynamics estimated using a lower Ct threshold will differ from observed dynamics if the distribution of Ct values is not constant over time (i.e., shift in viral load). Stronger shifts in Ct distribution will yield larger differences between observed and permuted datasets. (For further explanation, see Appendix S3). Permutations were done between samples of the same buffer type to account for bias introduced by preservation.

We compared prevalence dynamics estimated using original Ct values with 500 permuted prevalence curves. For this, we calculated the proportion of permuted prevalence values that were smaller than the observed prevalence value at each time-point (i.e., the observed prevalence quantile). If there was no change in the distribution of Ct values over time, the quantiles would be relatively constant around ∼50%. If Ct values shifted towards low values (high viral loads; e.g., during high-prevalence periods) and vice-versa, the quantiles would be high during high-prevalence periods and low during low-prevalence periods.

To visualize whether lower Ct thresholds improved the association between prevalence and spillover risk, we divided the time series by off-season months (September-May) (Eby et al. 2023) with no cases of spillover, and peak-season (winter; June-August) without and with cases of spillover, and plotted the prevalence estimated for each Ct threshold. We fitted generalized linear models (GLMs) to test (1) prevalence by time-series category across Ct values, and (2) prevalence by Ct value during peak season with spillover only (both fit with binomial error distribution and logit link function).

To test whether lower Ct thresholds improved the association between prevalence and spillover risk, we also fitted a GLM to the number of spillover events (Poisson error distribution and log link function) and the occurrence of spillover events (binomial error distribution and logit link function), for each Ct threshold. This was done for both the observed and the permuted prevalence estimates. A measure of model fit (AIC) was then used to assess: (1) whether model fit improved for lower thresholds, and (2) whether model fit improved relative to the permuted datasets. Analyses including spillover data were performed on sites aggregated across space (Appendix S1), and for the single study site (“Clunes”) spatially linked with spillovers during the duration of the study (Figure 1, Appendix S2).

## Results

### Description of primary dataset

We tested 6,151 unique pooled urine samples from 152 sampling sessions collected from July 2017 through September 2020 for the presence of Hendra virus (HeV-g1). For 4,657 of these samples, at least one black flying-fox was recorded above the collection sheet. Herein, we restrict analyses to samples associated with black flying-foxes as this is the primary reservoir for the prototypic Hendra virus variant tested. Using the conventional Ct 40 threshold, 361 of 4,657 samples were positive for Hendra virus (7.8%) with a mean viral load of 22,000 genome copies/ml.

### Virus isolation

In our primary dataset, virus was isolated from three samples with Ct values of 24, 35, and 37 (∼1,000,000 to 400 genome copies/ml), from a total set of 177 samples with broad range of Ct values (23-40) (Appendix S5). In the collated historical datasets, Hendra virus was isolated in 8 of 62 Hendra virus-positive samples collected from bats (with Ct values of 29, 34 and 36), horses (Ct: 27 and 29) and humans (Ct: 23) (Playford et al. 2010; Barr et al. 2015) (Appendix S5).

While overall isolation was low (an outcome typical for bat viruses) the relationship between isolation and Ct value suggested that infectious virus was more likely in samples with low Ct (Figure 2, Appendix S5). For every increase in Ct by 1, the probability of isolation decreased by 18.4% (±6.6%, p= 0.0014). At Ct 40 (the conventional threshold), virus could be isolated from only 1% of samples. This increased to 7.3% at Ct 30. This relationship was consistent when restricted to isolations from bat samples (8/223; 3.6%): for every increase in Ct by 1, the probability of isolation decreased by 17.6% (±8.4%, p= 0.0164). At Ct 40, virus could be isolated from only 0.98% of samples, and rose to 6.4% at Ct 30 (Appendix S5).

**Figure 2:**
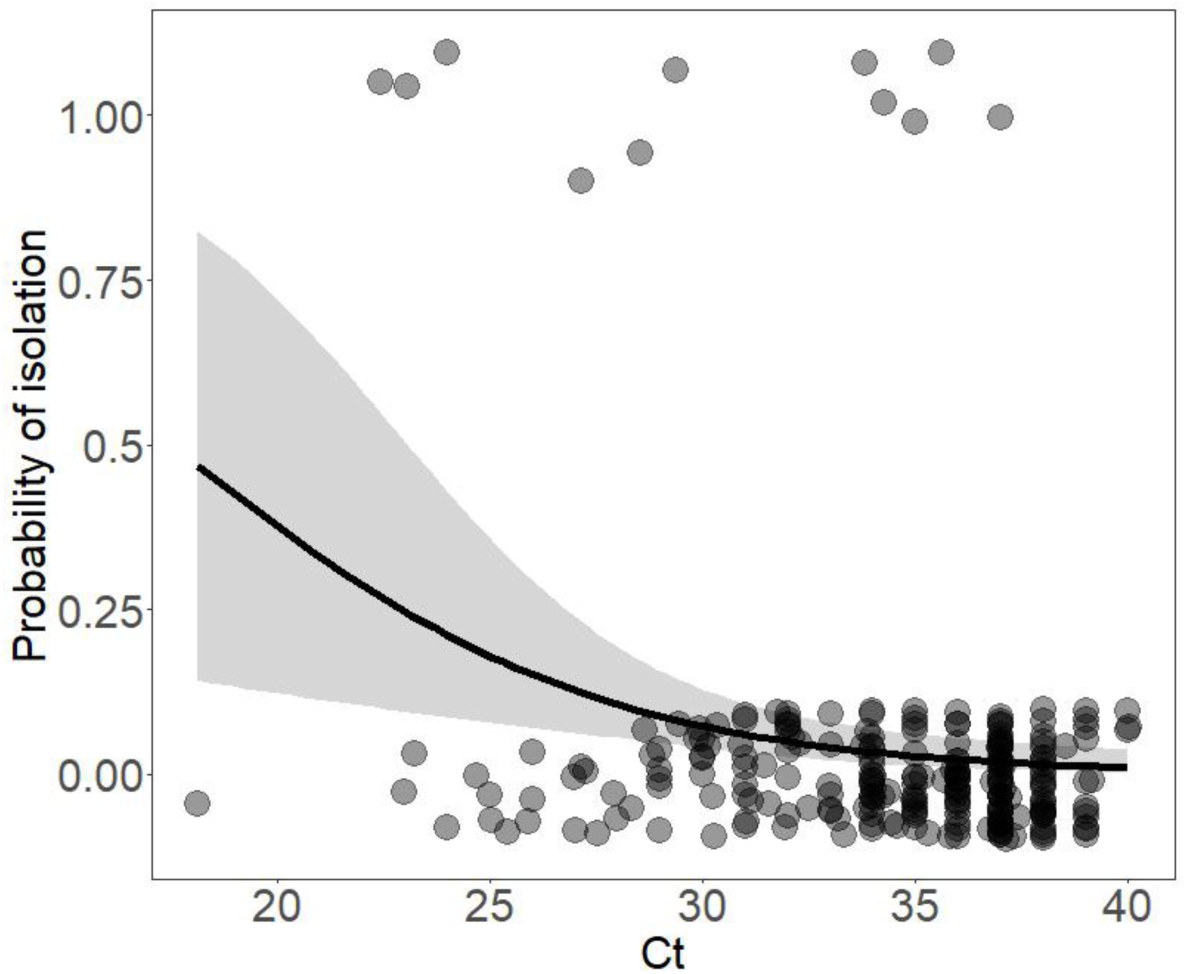
Logistic regression of isolation relative to Ct value of sample, from 239 Hendra positive samples taken from bats, horses and humans. A jitter is applied to the points to aid visualization, so that the plot shows successful (top) and unsuccessful (bottom) isolation attempts.

### Prevalence and viral load distribution

We observed seasonal and inter-annual patterns in shedding prevalence. With high Ct threshold values, viral excretion typically peaked in the winter peak season (Jun-Aug), but low-prevalence shedding was apparent throughout the rest of the year. Using lower thresholds, pooled prevalence was minimal outside of winter. Large peaks in prevalence were not observed every year, and this was consistent across all thresholds (Figure 3).

**Figure 3:**
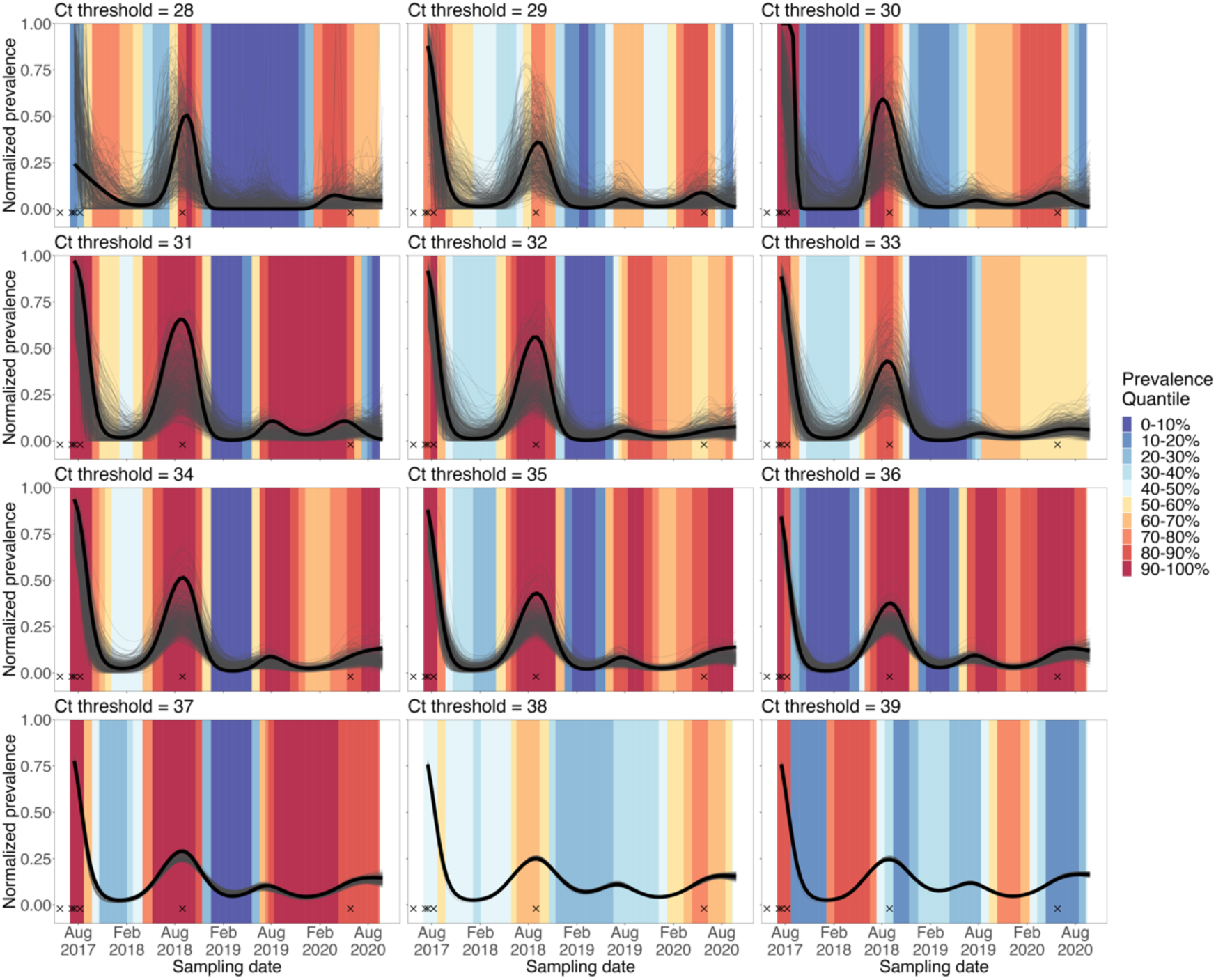
Permutation analysis at different Ct thresholds. For each Ct threshold, Ct values were permuted 500 times, and prevalence was re-estimated (thin grey lines). For each time-point, the quantile of observed prevalence (bold black lines) was calculated as the proportion of permuted prevalence values that was smaller than the observed value. Background colors indicate prevalence quantiles for each time-point. When Ct distributions are similar over time, prevalence quantiles are expected to be around 50%, whereas changing Ct distributions are expected to result in low and high quantiles. Prevalence is pooled for all sites and normalized to allow comparison between thresholds. Cases of spillover are shown with crosses. Note that a spillover occurred immediately prior to our sampling period (May 2017); this is included in the visual but not in analyses as there was no concurrent sampling.

Moreover, the permutation analysis showed strong statistical support for a Ct distribution shift towards disproportionately lower values (higher viral loads) during high-prevalence periods, and towards disproportionately higher values during low-prevalence periods (Figure 3). Across Ct thresholds, high quantiles coincided with high-prevalence (i.e., observed prevalence being higher than most permuted prevalence values) and low quantiles coincided with low-prevalence periods, but only for thresholds below Ct 38.

Cases of spillover clearly aligned with peaks in pooled prevalence estimated using lower Ct threshold values (Figure 3). When the time series was subdivided based on season and spillover event observation, prevalence was highest for peak seasons in which a spillover event was observed, and this was consistent for all Ct threshold values (Figure 4) (effect size: peak-season & spillover=13.7±1.43, peak-season & no spillover=2.33±1.34, p<0.0001, df =2, χ^2^=53.2). For peak season with spillover events, prevalence trended higher for lower Ct threshold values, excluding the two lowest values (Figure 4), although this was not statistically supported (effect size=0.90, p=0.30, df=1, χ^2^=1.12). These results were further supported by the permutation analyses, as the association between pooled prevalence and spillover was stronger for threshold values below Ct 38. This was shown by the low AIC quantiles for GLMs linking prevalence to number of spillover events, and to a lesser degree, for GLMs linking prevalence to occurrence of spillover events (Figure 5).

**Figure 4:**
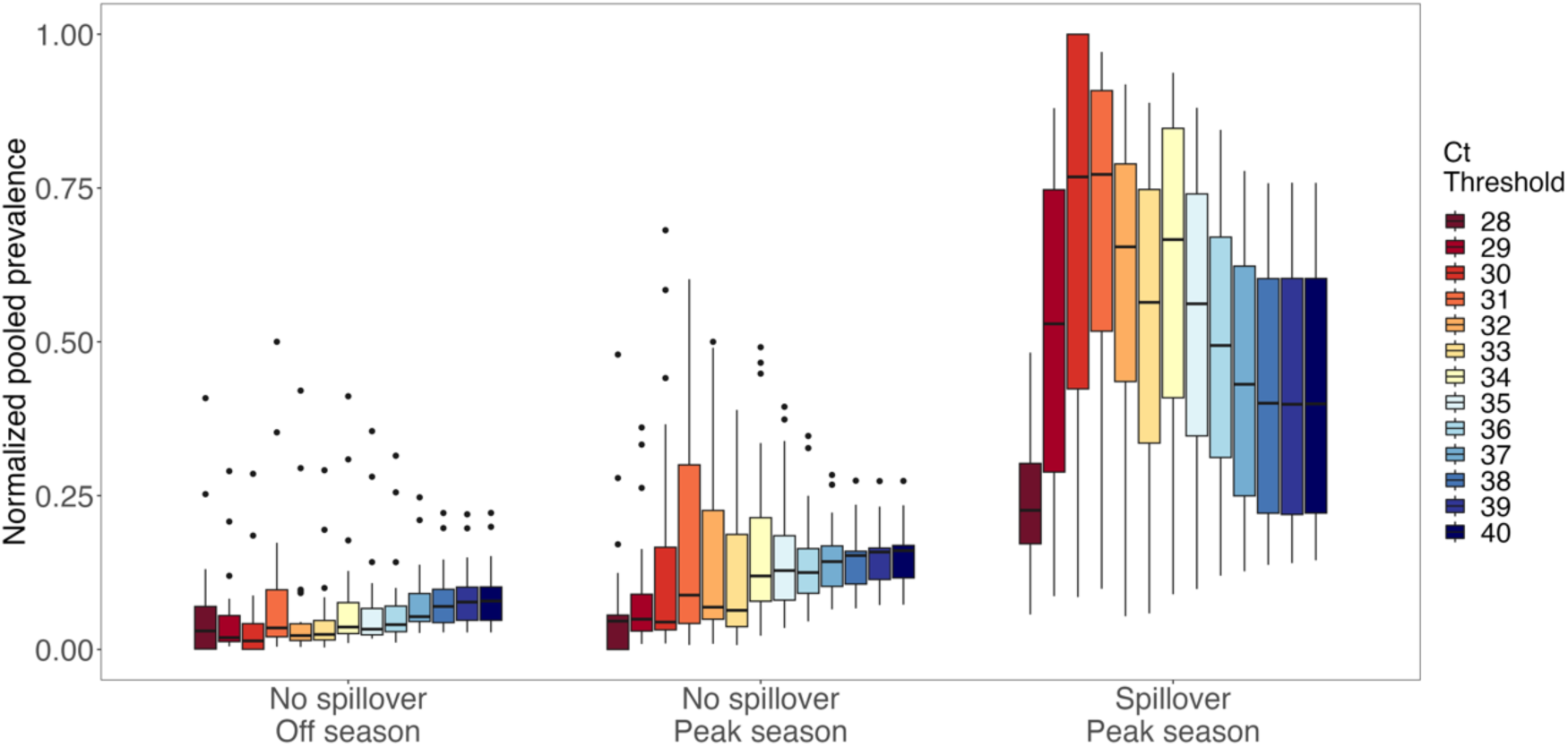
Normalized pooled prevalence for different Ct threshold values across off-season months (September-May) with no cases of spillover, and peak season (June-August) without and with cases of spillover. Boxplots show the 25%, 50%, 75% percentiles, lines indicate the smallest and largest values within 1.5 times the interquartile range, dots indicate values beyond that. Normalization is intended to show the relative changes in prevalence across the time series (therefore, how the dynamics of prevalence varies), for each Ct threshold. For non-normalized prevalence values at each Ct threshold, see Appendix S1.

**Figure 5:**
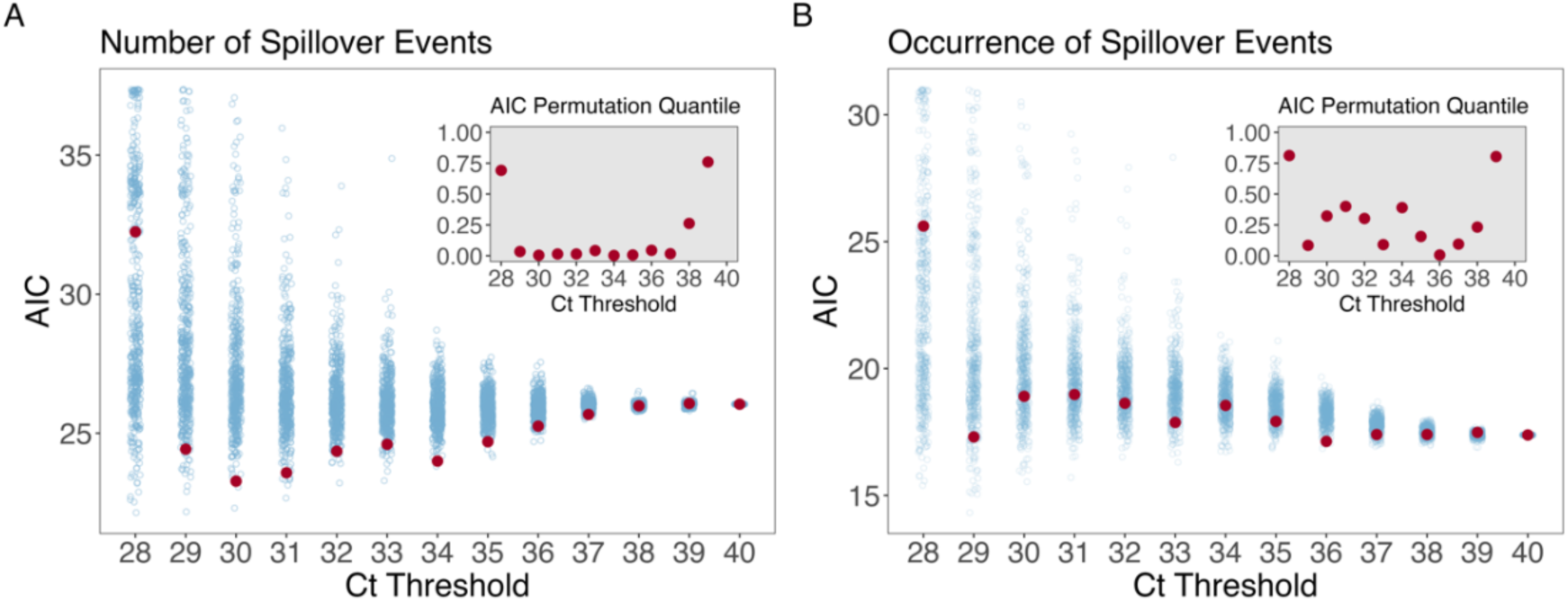
Akaike Information Criterion (AIC) values for models with normalized pooled prevalence as predictor variable and number (A) or occurrence (B) of spillover events as outcome variable, for a range of Ct thresholds. AIC values are shown in blue for permuted data and red for observed data. Insets show the proportion of permuted AIC values that is larger than the observed value. Low quantile values provide statistical support for a shift in the distribution of Ct values that results in a better correlation between prevalence and spillover.

## Discussion

We give evidence of periodic shifts in viral load distribution in bat populations, skewing toward higher viral loads of Hendra virus during high-prevalence periods. These shifts lead to a disproportionate increase in risk of spillover due to the combination of higher prevalence and higher viral loads being shed. Despite the small number of spillover events during our study period, there was statistical support for a correlation between viral load distributional shifts and spillover. Additionally, using more conservative Ct thresholds generated prevalence estimates that more strongly correlated with spillover risk. Collectively, these findings suggest that conventional detection of Hendra virus RNA at Ct 40 may not be the best for predicting spillover.

The Hendra virus RT-PCR (TaqMan) assay was developed to diagnose Hendra virus (genotype-1) infection specifically and rapidly, to allow prompt implementation of control measures on infected properties (Daniels et al. 2001; Smith et al. 2001). The priority—to identify infected but not necessarily infectious animals—balances the need for a high detection rate (where the consequences of a missed assessment are great) with a small proportion of false positive results (Daniels et al. 2001). In clinical settings where a failure to act may result in delayed treatment and human fatality, a high probability of detection with few misses is appropriate. Detection of Hendra virus RNA for exclusion testing uses a Ct 40 threshold for this reason (e.g., Field et al. 2010). Wildlife epidemiological studies, by contrast, pre-emptively identify spatiotemporal patterns of risk to inform future management. Modelling tools often assume that RNA detection equates to infectiousness (i.e., susceptible, infectious, recovered models) (e.g., Plowright et al. 2016). In these scenarios, identification of hosts that are actively infectious is an appropriate priority. Though the precise functional relationship between infectivity and viral load is currently unknown for even the most well studied viruses (e.g., SARS-CoV-2), it is widely accepted that individuals with higher viral loads are more likely to be infectious (Hay et al. 2021; Snedden and Lloyd-Smith 2023).

While lower Ct thresholds generally improved the association between prevalence and spillover, we found no specific optimal value, nor a clear trend towards stronger correlations for lower values. This may be due to statistical limitations, as lower thresholds create smaller sample sizes which decrease statistical power. It is also possible there is no difference among thresholds below Ct 38. This could be the case if higher thresholds, such as the conventional Ct 40 threshold, do not adequately differentiate infectious and non-infectious samples, while lower thresholds include proportionally more (or exclusively) infectious samples. It is not clear from the isolation–Ct correlation which explanation is more likely, as we see that isolation can be successful for Ct≤37, and – although the fitted model suggests a negative correlation – the confidence intervals are wide.

Using lower thresholds, pooled prevalence was minimal outside of peak season, and large peaks of infectiousness were not observed every year. Additionally, normalized prevalence estimated using lower thresholds was higher than the conventional estimate during peak seasons where spillover events were observed, but lower during peak seasons where no spillover events were observed, or during the off season. Collectively, this suggests that prolonged low-intensity shedding of Hendra virus outside of winter months – captured only with the conventional threshold – may reflect prolonged shedding of non-infectious RNA, or viral loads that are insufficient or unlikely to overcome dose barriers to spillover infection. More broadly, this association may impact relative spillover risk of Hendra virus variants, given the low prevalence and low viral loads of Hendra virus genotype-2 in wild flying fox populations, relative to genotype-1 (Peel et al., 2022)

Various mechanisms have been proposed to explain patterns in Hendra virus infection in subtropical Australia, including the temporal availability of eucalyptus blossom – the preferred food of flying-foxes (Parry-Jones and Augee 1991; Kessler et al. 2018; Eby et al. 2023). It has been proposed that low food availability and utilization of alternative diet sources may lead to increased viral shedding via reduced immunocompetence and reactivated infection (Plowright et al. 2016; Becker et al. 2023). In winter, few species of eucalyptus reliably produce nectar (House 1997; Eby and Law 2008); this period of food scarcity coincides with the higher levels of Hendra virus shedding in subtropical regions reported here and elsewhere (Field et al. 2015; Páez et al. 2017). Further analyses will be required to assess whether food availability and nutritional composition explains distributional shifts in viral load.

To further explore the relationship between viral load proxies and infectiousness, future studies could formally test a serial dilution relationship between RNA detection (genome copy numbers) and *in vitro* infectiousness (Median Tissue Culture Infectious Dose, TCID_50_) of henipaviruses on different cell lines (Munster et al. 2009; Bullard et al. 2020). As this relationship becomes clearer, future models of bat-pathogen transmission dynamics may also consider RNA shedding to incorporate within-host infection dynamics into spillover risk estimates. Future studies that seek to compare Ct values with results in this study should consider that Ct values alone are not directly comparable across laboratories (Munster et al. 2009).

## Conclusion

The intensity of viral shedding is a valuable but often overlooked component of wildlife pathogen surveillance. For viruses where RNA shedding can be detected long after disappearance of infectious virus, estimation of prevalence without consideration of viral load can decouple the presence of virus from the risk of transmission. We suggest that circulation of Hendra virus in bat populations – quantified conventionally as prevalence of viral RNA at Ct≤40 – may not be sufficient for predicting spillover risk. We show that prevalence estimated with more conservative threshold values can provide a more informative measure that better aligns with Hendra virus spillover risk. Moreover, we suggest that previously reported low-intensity shedding does not align with spillover risk and may not reflect excretion of infectious virus. By integrating information on viral load proxies with estimates of prevalence, our results refine predictions of spillover risk and will contribute to the future management of disease risk from this virus. We propose that consideration of viral load (or viral load proxies) along with prevalence could improve inferences based on longitudinal surveys of wildlife viruses.

## Acknowledgements

We acknowledge the Kabi Kabi, Turrbal, Widjabul Wia-bal, Yugambeh and Yuggera Ugarapul people, who are the Traditional Custodians of the land upon which this work was conducted. We would like to thank Tim Pearson, Denise Karkkainen, Ticha Padgett-Stewart, Justine Scaccia, Ariana Ananda, Emma Glennon, Kimberley Martyn, Rhiannon Kirton, Hannah Eiseman, Michael Johnson, Kathryn Yock, Eloise Skinner, Jackelyn Urquhart, Renata Muylaert, Jaylan Shabrod, Alice Risely, Ashleigh Murray, Vee Halilovic, Lee-anne Lunn, Kate Dutton-Registor, Kurt Winter, Melissa Walker, Geoffrey Smith, Cinthia Pietromonaco, Laura Pulscher, Cecilia Sanchez, Kerryn Perry-Jones, Cara Parsons, Jessica Brito, and Julia Strazz for their assistance in the field. We also thank Isaac Knights, Dian Riseley, and Stella Maris Januario da Silva for project support, and the Parks family and other landholders for kindly granting us access to property. This research was conducted under a Griffith University Animal Research Authority permit (DEB-1716698), a Scientific Purposes Permit from the Queensland Department of Environment and Heritage Protection (WISP17455716), a permit to Take, Use, Keep or Interfere with Cultural or Natural Resources (Scientific Purpose) from the Department of National Parks, Sport and Racing (WITK18590417), a Scientific License from the New South Wales Parks and Wildlife Service (SL101800) and general and products liability protection permit (GRI 18 GPL), and with permission to undertake research on council and private land.

