## Supplemental Information for "Periodic shifts in viral load increase risk of spillover from bats"

### Appendix S1: Spatiotemporal patterns in (not normalized) prevalence

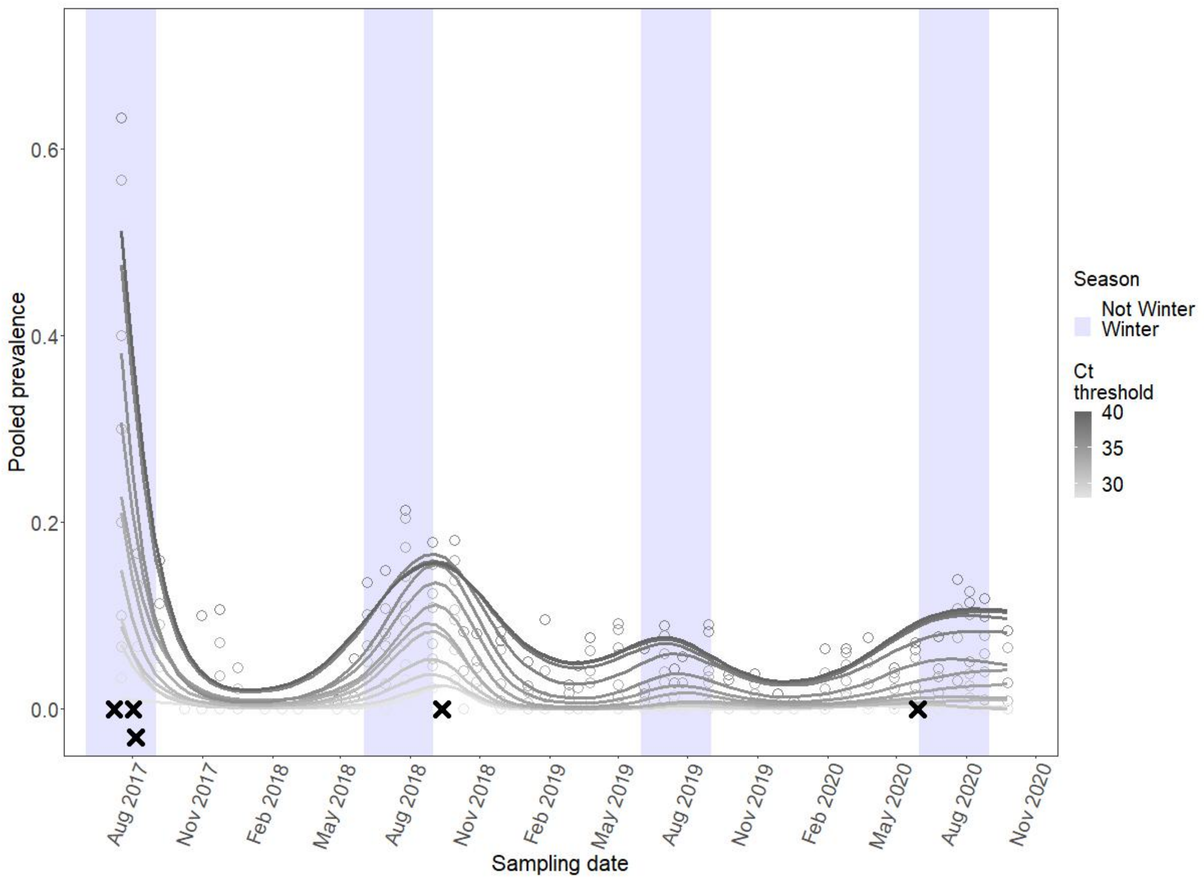


Figure S1: Predicted pooled prevalence (not normalized) from GAMs fit to the time-series site-aggregated data, estimated using different Ct thresholds. Cases of spillover are shown with crosses.

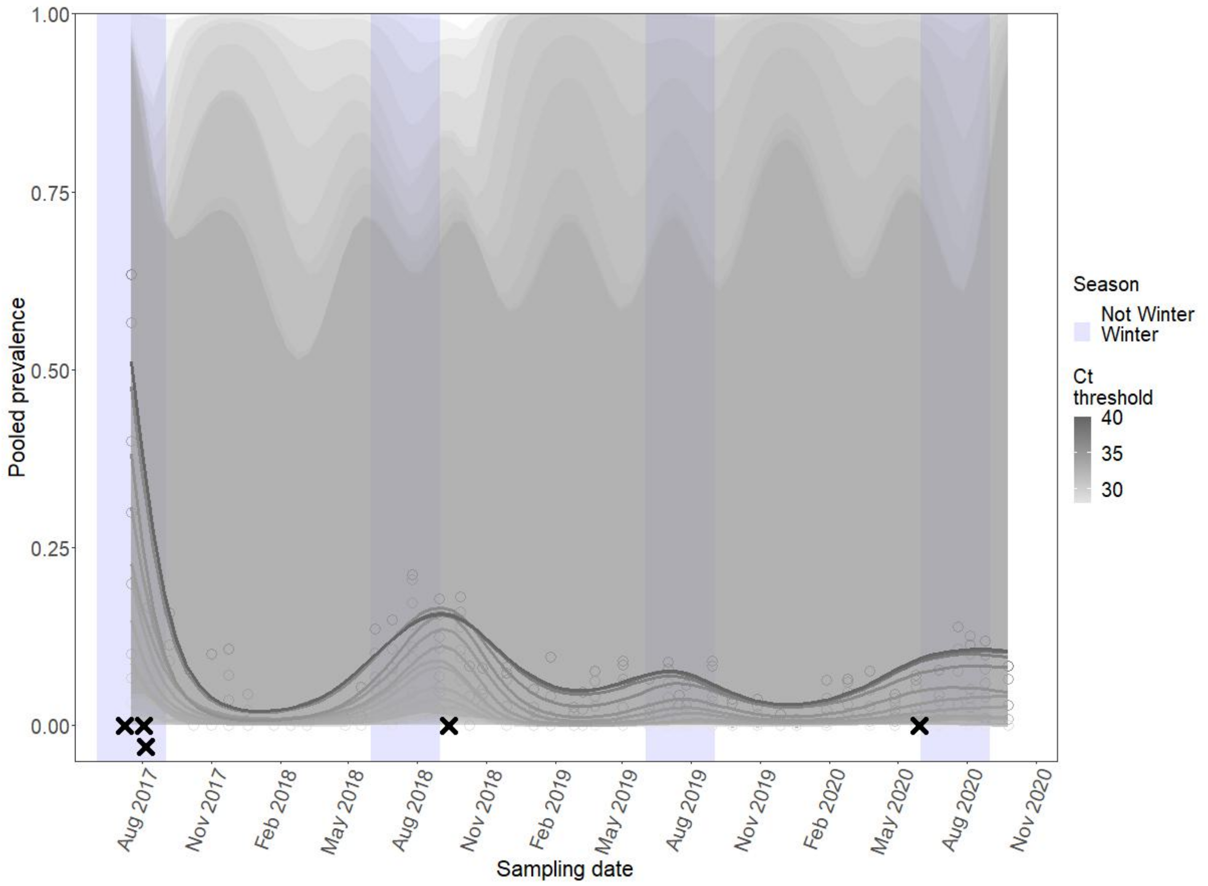


Figure S2: Predicted pooled prevalence (not normalized) with standard error, from GAMs fit to the time-series site-aggregated data, estimated using different Ct thresholds. Cases of spillover are shown with crosses.

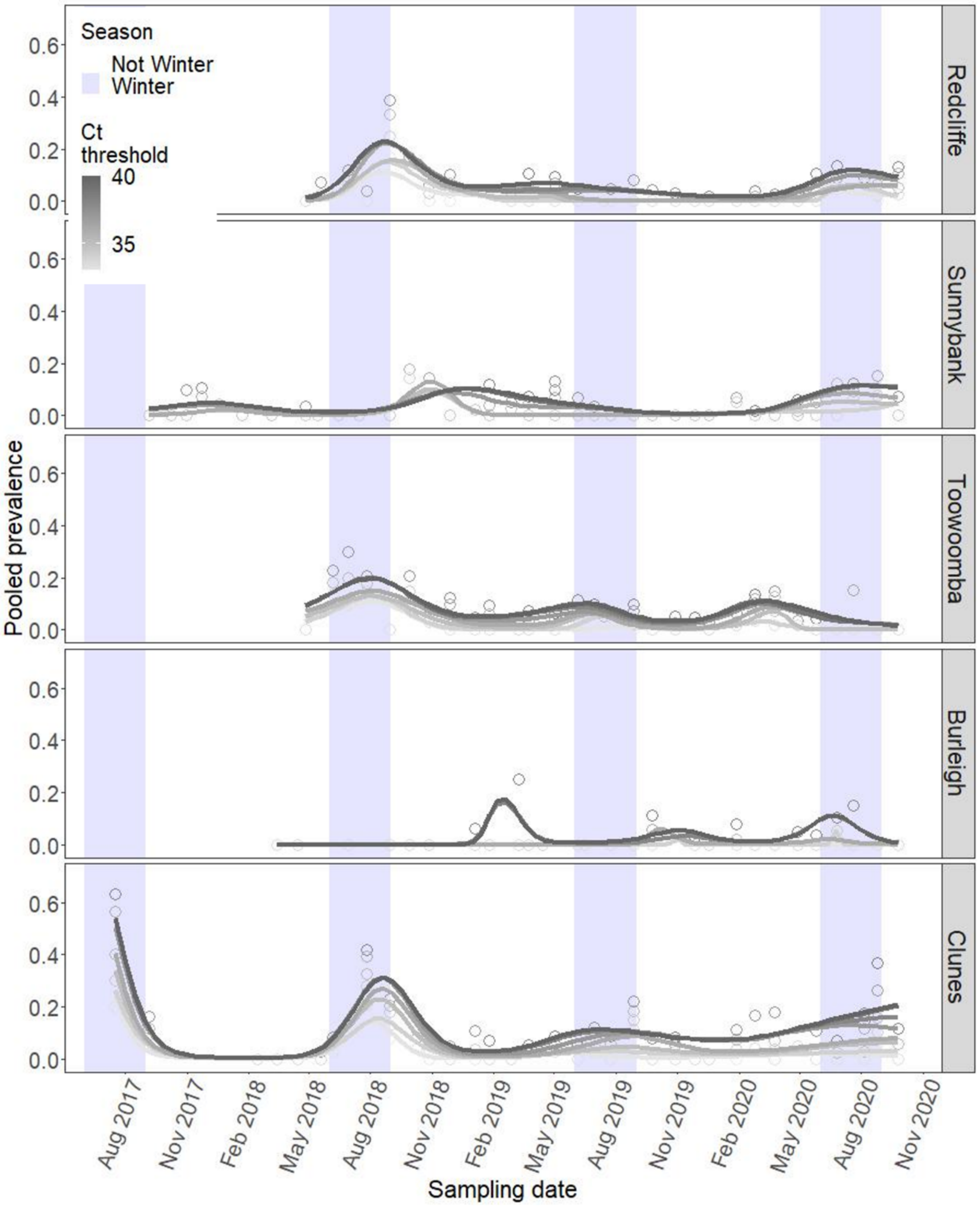


Figure S3: Predicted pooled prevalence (not normalized) from GAMs fit to the time-series data, per site, estimated using different Ct thresholds.

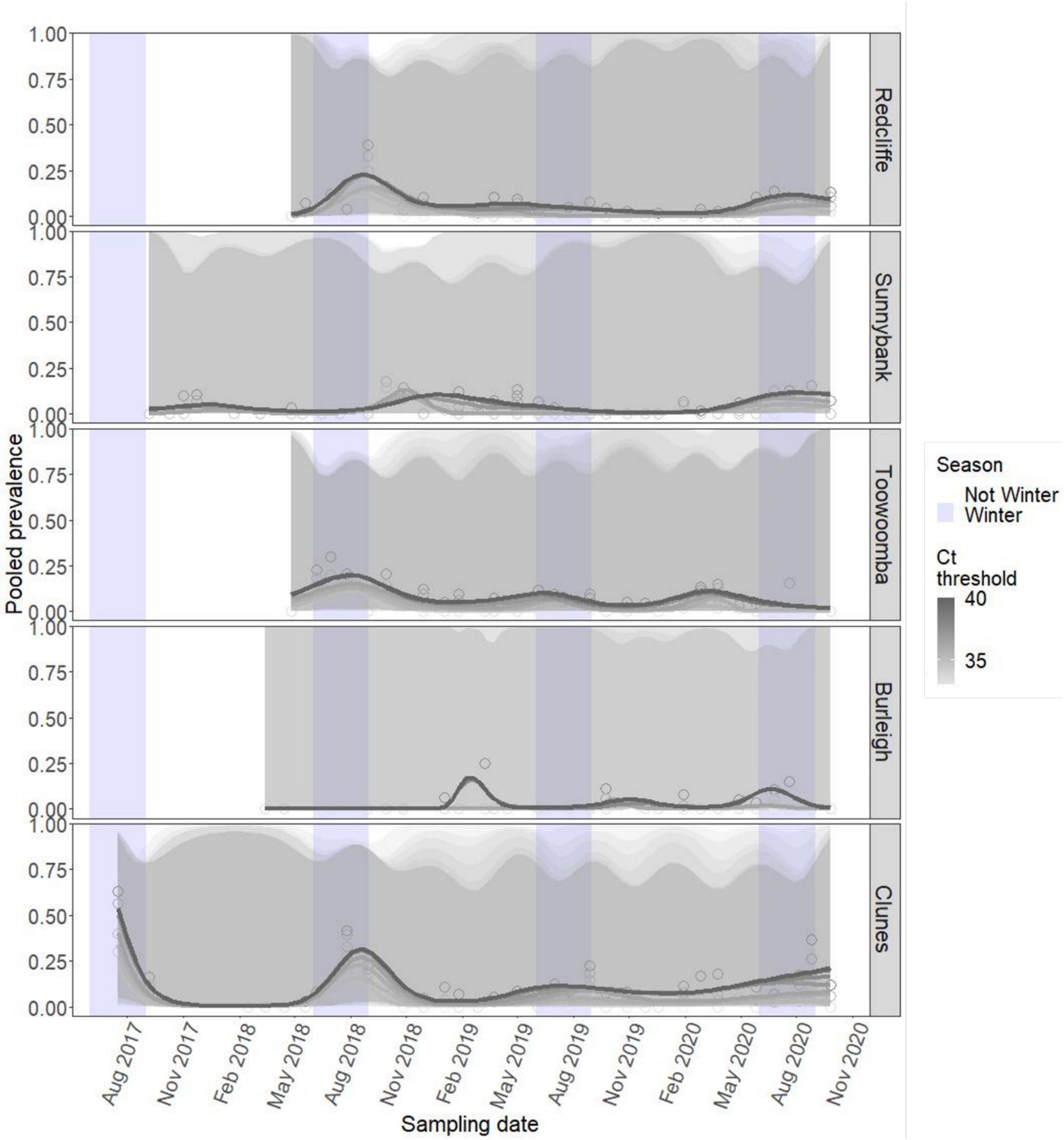


Figure S4: Predicted pooled prevalence (not normalized) with standard error, from GAMs fit to the time-series data, per site, estimated using different Ct thresholds.

**Appendix S2: Analyses of a single site (Clunes)**

Analyses including spillover data were performed on sites aggregated across space, and for the single study site ("Clunes") that was spatially linked with spillovers during the duration of the study. In addition to having key features associated with higher risk of spillover (continuous occupation by black flying-foxes, *Pteropus alecto*; recently established as overwintering; and highly restricted access to native winter food sources), this site, "Clunes", showed high use of agricultural areas by bats (Figure S5). This attribute has also been linked with spillover occurrence, because horses may be present at higher densities in agricultural areas than in urban and forested areas (Eby et al. 2023).

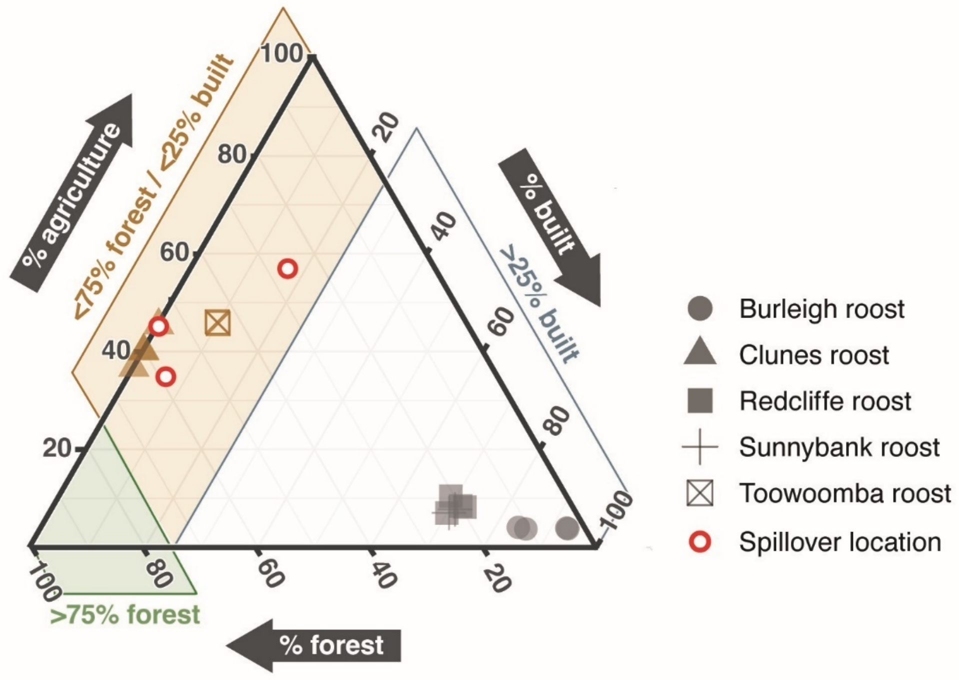


Figure S5: Proportion of foraging area surrounding study roosts that was classified as built, forested or agricultural, per study year (2017-2020) (Eby et al. 2023). The red circles indicate roosts that were the sources of winter spillovers.

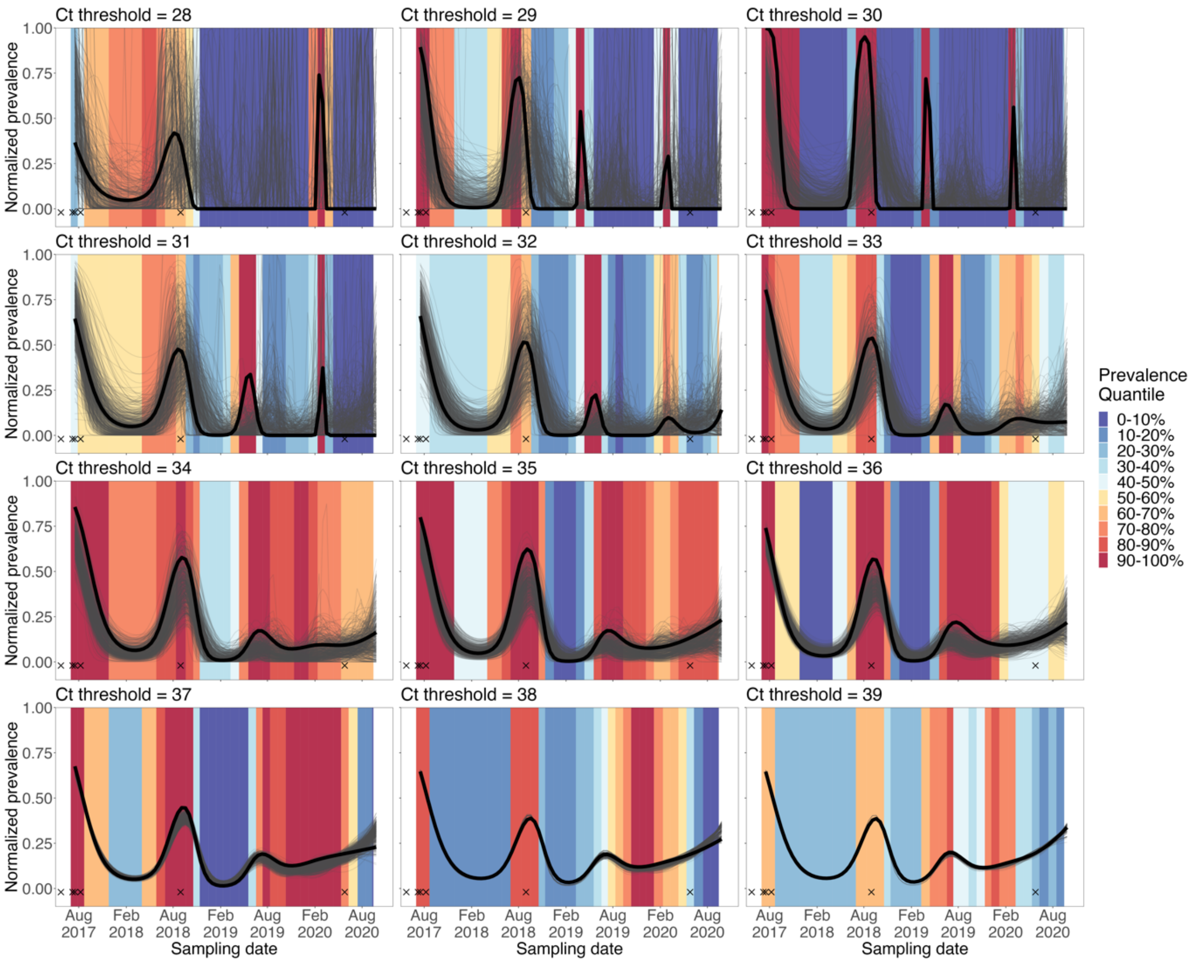


Figure S6: Permutation analysis at different Ct thresholds **for Clunes**. For each Ct threshold, Ct values were permuted 500 times, and prevalence was re-estimated (thin grey lines). For each sampling session, the quantile of observed prevalence (bold black lines) was calculated as the proportion of permuted prevalence values that was smaller than the observed value. Background colors indicate prevalence quantiles for each session. When Ct distributions are similar over time, prevalence quantiles are expected to be around 50%, whereas changing Ct distributions are expected to result in low and high quantiles. Prevalence is pooled for all sites and normalized to allow comparison between thresholds. Cases of spillover are shown with crosses.

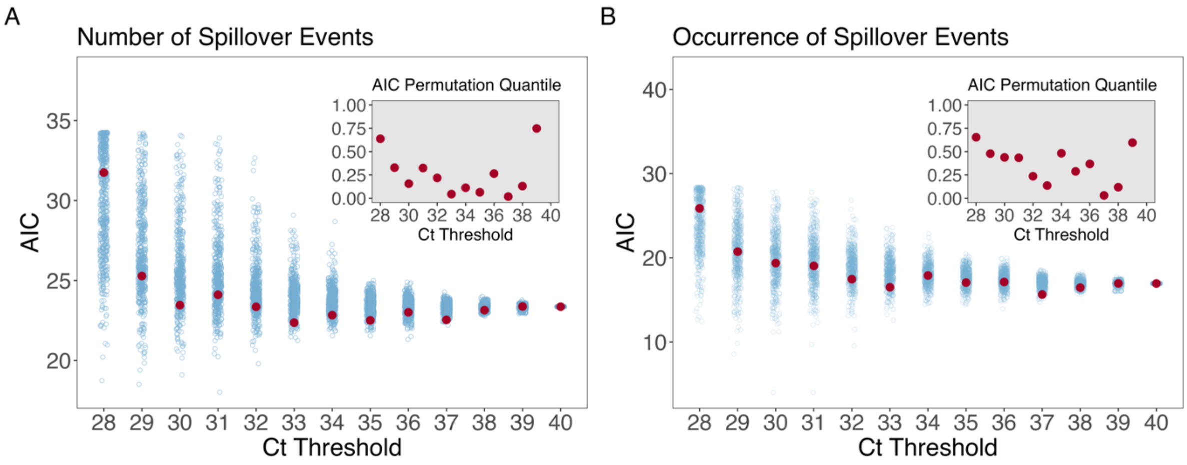


Figure S7: Akaike Information Criterion (AIC) values for models with normalized prevalence **at Clunes** as predictor variable and number (A) or occurrence (B) of spillover events as outcome variable, for a range of Ct thresholds. AIC values are shown in blue for permuted data and red for observed data. Insets show the proportion of permuted AIC values that is larger than the observed value. Generalized linear models with Poisson error distribution and log link function (number of spillover events) and binomial error distribution and logit link function (occurrence of spillover events) were used. Low quantile values provide statistical support for a shift in the distribution of Ct values that results in a better correlation between prevalence and spillover.

### Appendix S3: Supporting materials and methods

*Site selection*

We conducted our study at five roosts spanning south-east Queensland to north-east New South Wales. Roosts were selected because they have attributes that are associated with higher risk of spillover: 1) continuous occupation by black flying-foxes (*Pteropus alecto*); 2) recently established as overwintering; and 3) highly restricted access to native winter food sources (Eby et al. 2023). In addition, the roost population was consistently large enough for reliable sample collection (total estimate consistently >500); had suitable access and sampling conditions; was not a site of high human-bat conflict; and permissions could be obtained for long-term sampling.

*Sample collection*

Our "under roost" sampling approach followed an optimized sampling design described by Giles et al. (2018), to allow for better estimation of true prevalence with minimum sampling bias and false negatives, compared with under roost sampling methods used in other large scale sampling efforts of Hendra virus (Field et al. 2011; Field et al. 2015; Field et al. 2015; Burroughs et al. 2016). Key differences in our methodology included: 1) collection of a single, pooled sample per sheet, compared with 3-4 pooled samples per sheet used previously, 2) spacing of sheets at least 1 meter apart to ensure independence of samples, 3) collection from a higher number of smaller sheets: an average of 40, 0.9 x 1.3 meter sheets compared with 10, 3.6 m x 2.6 m sheets in e.g., Field et al. (2015), and 4) recording of the number and species overhead that likely contributed to the pooled sample.

*qRT-PCR screening of Hendra virus*

Table S1: Primer and probe sets used for Hendra virus screening. Primer and probes were provided by the Public Health Agency of Canada

| **Screening primers** | |
|---|---|
| **Hendra virus** | |
| DRP064- Hendra_F | CCCAACCAAGAAAGCAAGAG |
| DRP065-Hendra_R | TTCATTCCTCGTGACAGCAC |
| DRP66-Hendra_VIC | **VIC-**TTACTGCGGAGAATGTCCAACTGAGTG-**QSY** |

Table S2: Cycling program for Hendra virus screening

| **Step** | **Temperature** | **Time** | **Cycles** |
|---|---|---|---|
| Reverse transcription | 50°C | 5 minutes | 1 |
| RT inactivation / initial denaturation | 95°C | 20 seconds | 1 |
| Denature | 95°C | 15 seconds | 40 |
| Anneal / extend | 60°C | 60 seconds | 40 |

Table S3: Conversion between Ct value and genome copies per mL. Note that this conversion table is not universal across assays, and cannot be applied to assays from another laboratory.

| Threshold | Ct value | Mean genome copies per mL |
|---|---|---|
| Conventional | 40 | 59 |
| | 39 | 110 |
| | 38 | 203 |
| | 37 | 377 |
| | 36 | 699 |
| | 35 | 1297 |
| | 34 | 2404 |
| | 33 | 4459 |
| | 32 | 8268 |
| | 31 | 15332 |
| | 30 | 28431 |
| | 29 | 52722 |
| | 28 | 97766 |
| | 27 | 181297 |
| | 26 | 336194 |
| | 25 | 623433 |
| | 24 | 1156084 |
| | 23 | 2143826 |
| | 22 | 3975479 |
| | 21 | 7372070 |

We tested a total of 6,151 unique pooled urine samples from 152 sampling sessions collected from July 2017 through September 2020 for the presence of Hendra virus. At least one black flying-fox was recorded above the sheet of 4,657 samples. The number of samples collected per sampling session varied by month and site, depending on the roost extent and population size (and so, availability of space for sampling sheets). An average of 40 total samples were collected per sampling session (interquartile range: 31-50), with 32 on average (23-41) with black flying-foxes recorded above the collection sheet.

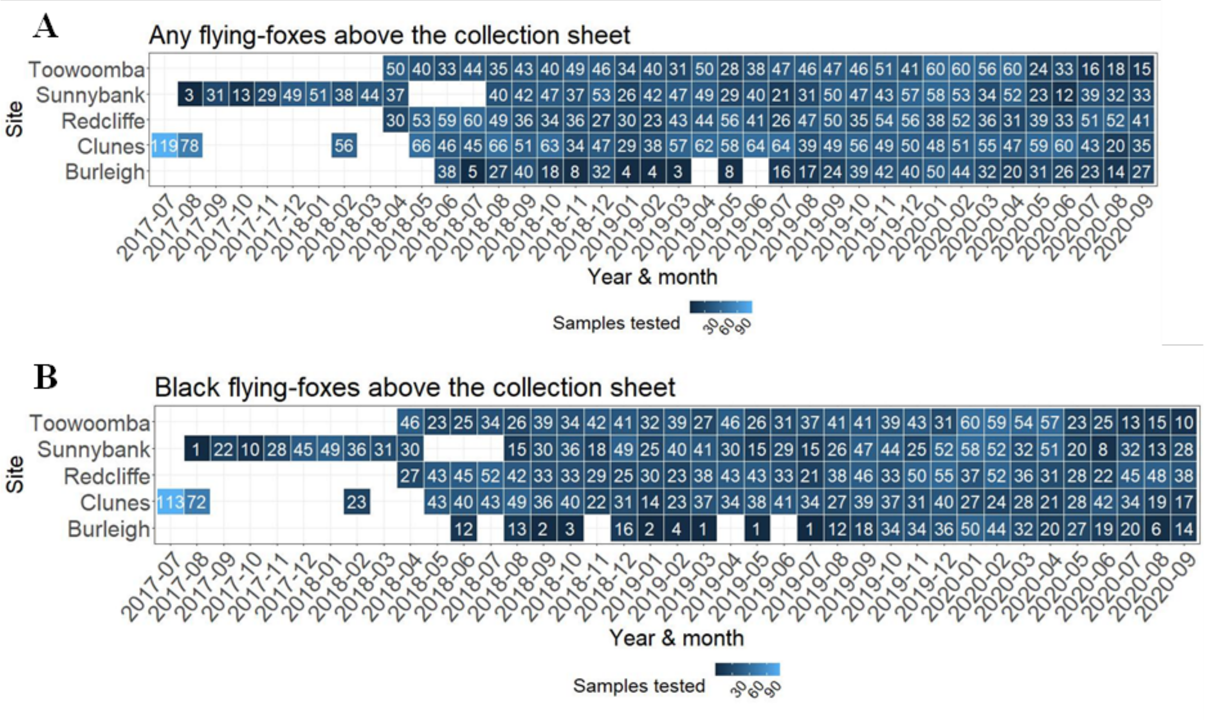


Figure S8: Number of samples taken per site over time. (A) Shows total samples, (B) shows samples with black flying-foxes recorded above the collection sheet.

*Permutation analysis for analyzing viral load distribution*

We assessed whether viral load follows a constant population distribution, or whether there are periods during which proportionally more individuals shed high viral loads. The former scenario would predict spillover risk to increase during peak periods because a larger number of individuals are shedding virus, and the latter scenario for spillover risk to increase because there are both more shedding individuals and more individuals that shed high viral loads. The latter would further predict spillover risk during low-prevalence periods to be disproportionately low, due to the lower proportion of individuals shedding high viral loads.

Permutations randomly shuffled Ct values of positive samples to re-estimate prevalence dynamics with randomized time. Prevalence dynamics estimated using a lower Ct threshold will differ from observed dynamics only if the population distribution of Ct values is not constant over time (i.e., a shift in viral load). Stronger shifts in Ct distribution will yield larger differences between observed and permuted datasets.

To explain: consider a scenario in which the distribution of Ct values shifts toward lower values (i.e., higher viral loads) during periods of high prevalence, with a corresponding shift toward high values during periods of low prevalence. Applying a lower Ct threshold to such a dataset would result in relatively more samples remaining positive during high-prevalence periods than during low-prevalence periods, and therefore relatively higher peaks and lower troughs. However, if the Ct values of positive individuals are first shuffled randomly (permuted), the same proportion of samples is expected to remain positive regardless of whether prevalence is high or low.

**Appendix S4: Effects of methodological choices on Hendra virus detection**

In this appendix we present visuals to check the effects of methodological choices and environmental influences on Hendra virus detection. Specifically, we tested for the effects of: time of sample collection, sample volume, temperature at the time of sample collection, whether samples had begun to evaporate or were in direct sun by the time of sample collection, and sample buffer (AVL, VTM or none). These checks were done to ensure sample quality prior to the formal analysis.

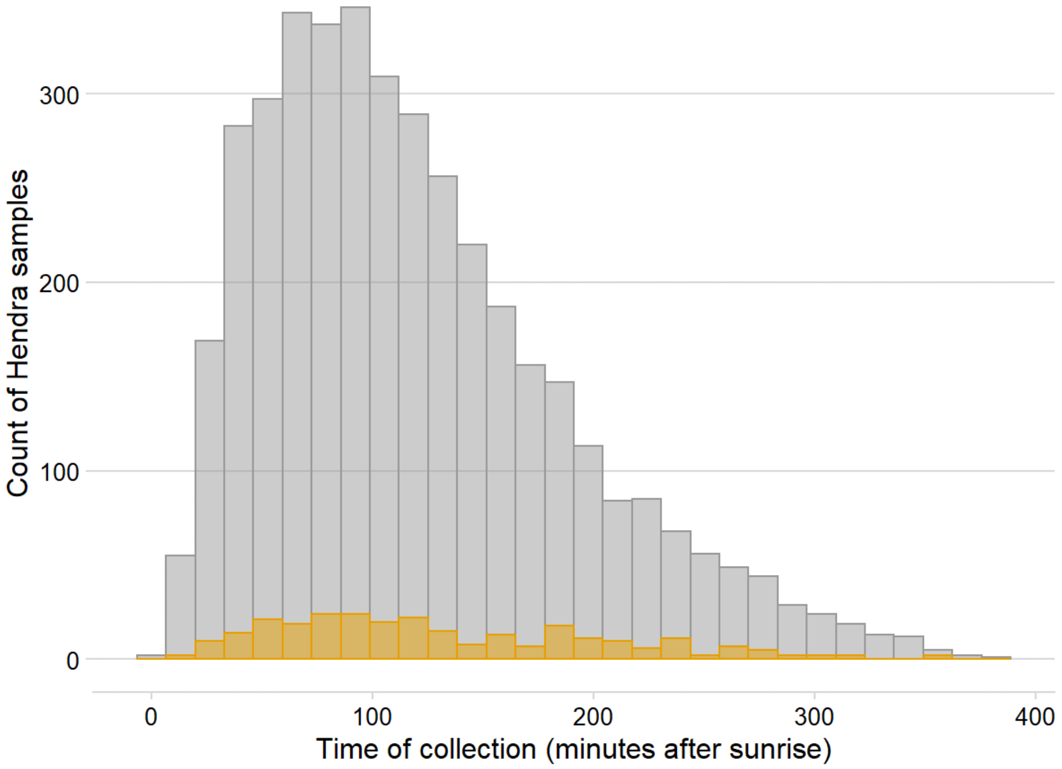


Figure S9: Time of sample collection and Hendra virus detection for continuously sampled Queensland roost sites. Grey bars indicate samples negative for Hendra virus, and yellow bars samples positive for Hendra virus. There is no systematic relationship between time of collection and number of Hendra virus detections. Note that time of collection was recorded for only a subset of samples.

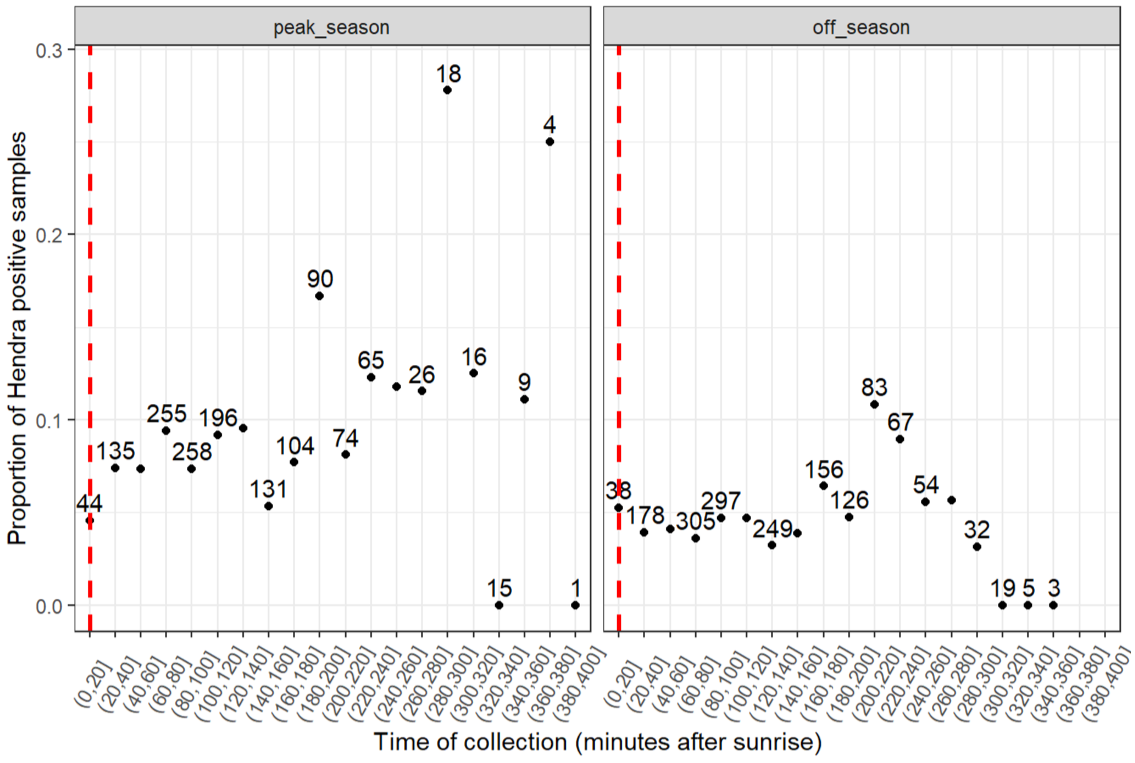


Figure S10: Time of sample collection and proportion of Hendra virus detections for continuously sampled Queensland roost sites. Facets divide samples collected into peak-season (June-August; time of peak Hendra virus shedding and detection) and off-season months. For periods with higher numbers of samples (>20 samples, number labels) there is no systematic relationship between time of collection and proportion of Hendra virus detections. Note that time of collection was recorded for only a subset of samples.

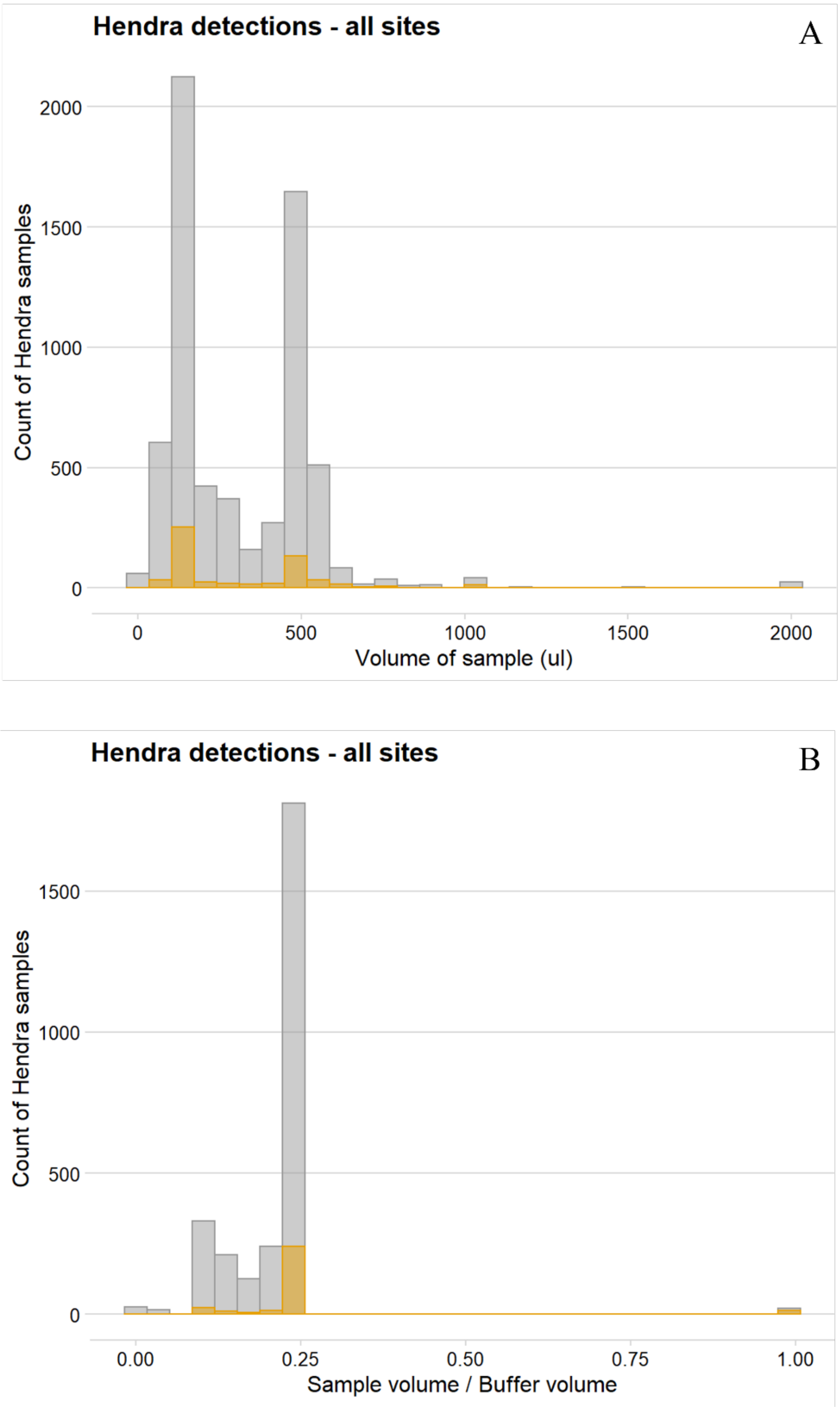


Figure S11: Volume of sample (A) and ratio of sample to buffer (B) and Hendra virus detection, for all roost sites. There is no systematic relationship between sample volume and number of Hendra virus detections. Note that exact urine volume was recorded for only a subset of samples.

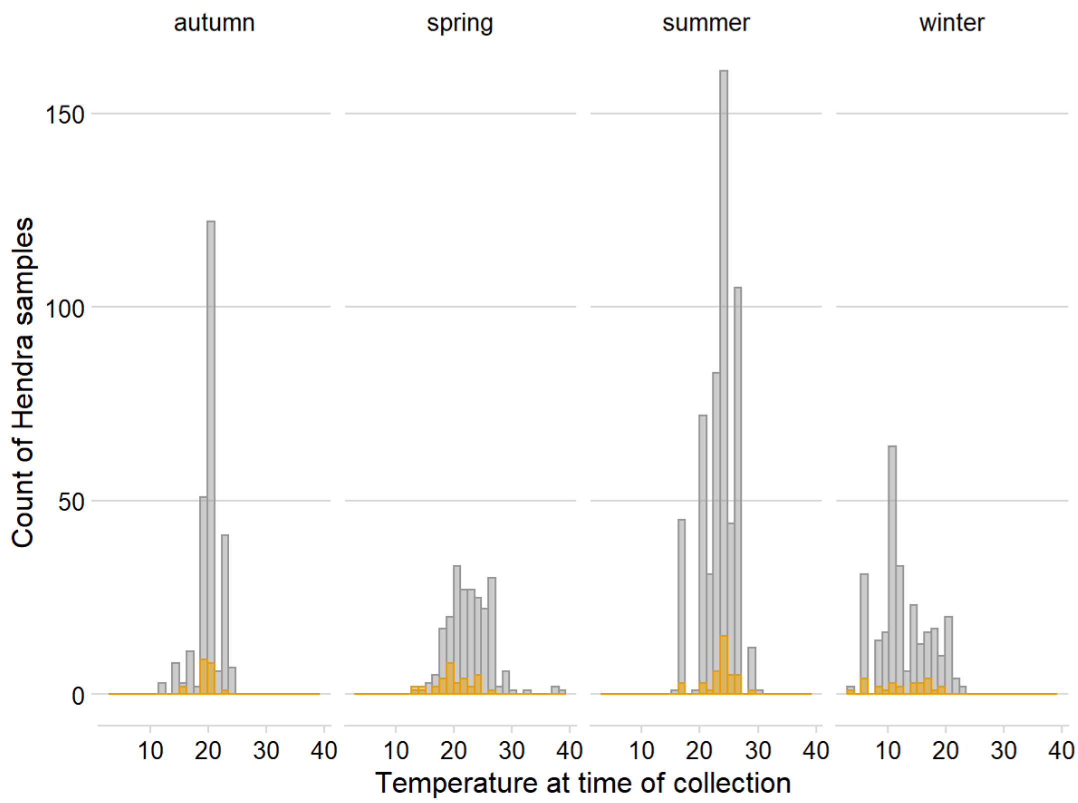


Figure S12: Temperature at time of sample collection and Hendra virus detection, for all roost sites. There is no systematic relationship between temperature at time of collection and number of Hendra virus detections within seasons. Note that temperature at time of collection was recorded for only a subset of samples.

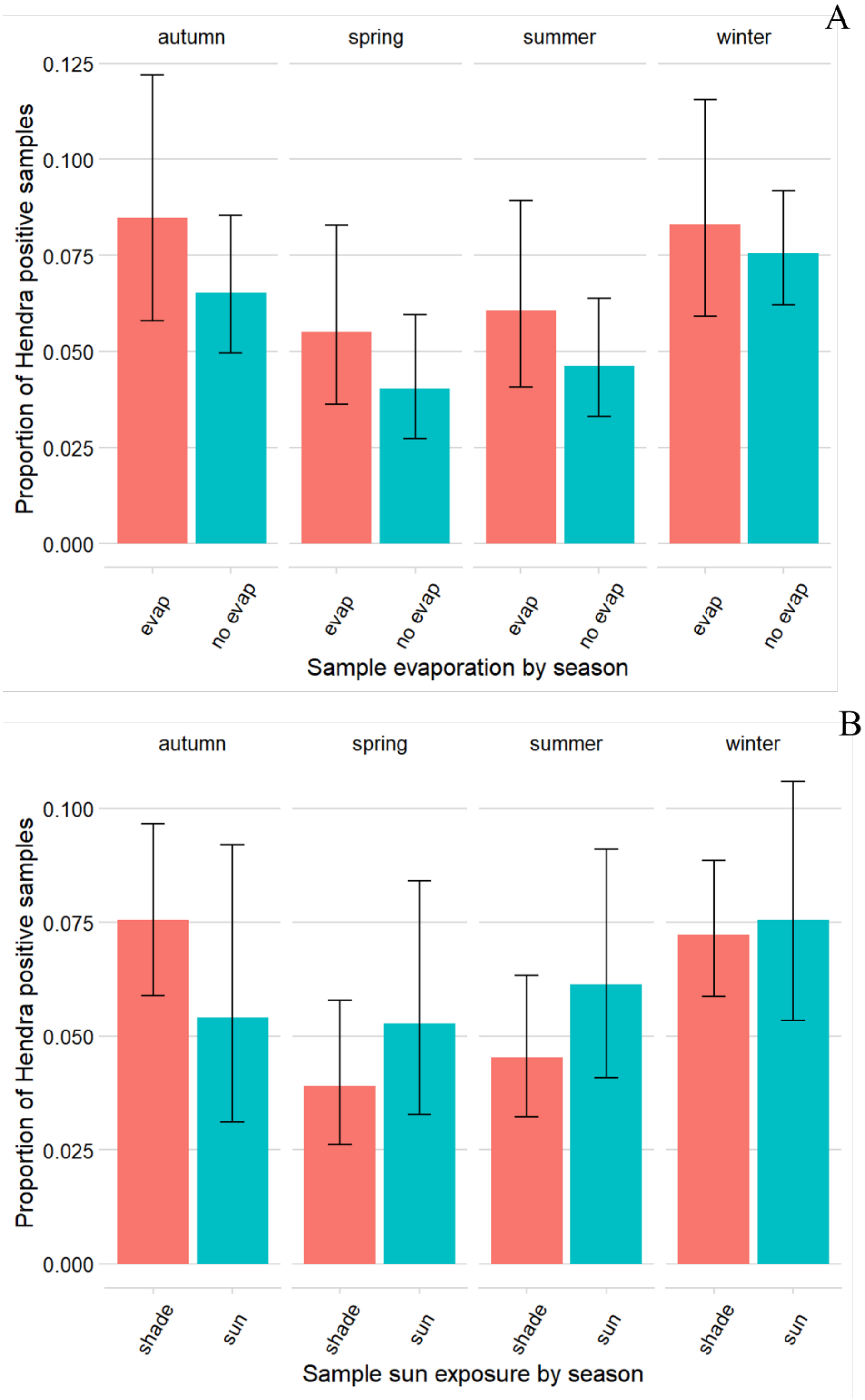


Figure S13: Effect of sun exposure (A) and evaporation (B) at time of sample collection on the proportion of Hendra positive samples. There is no systematic relationship between sun exposure and sample evaporation on Hendra virus detection. Note that evaporation and sun exposure information was recorded for only a subset of samples.

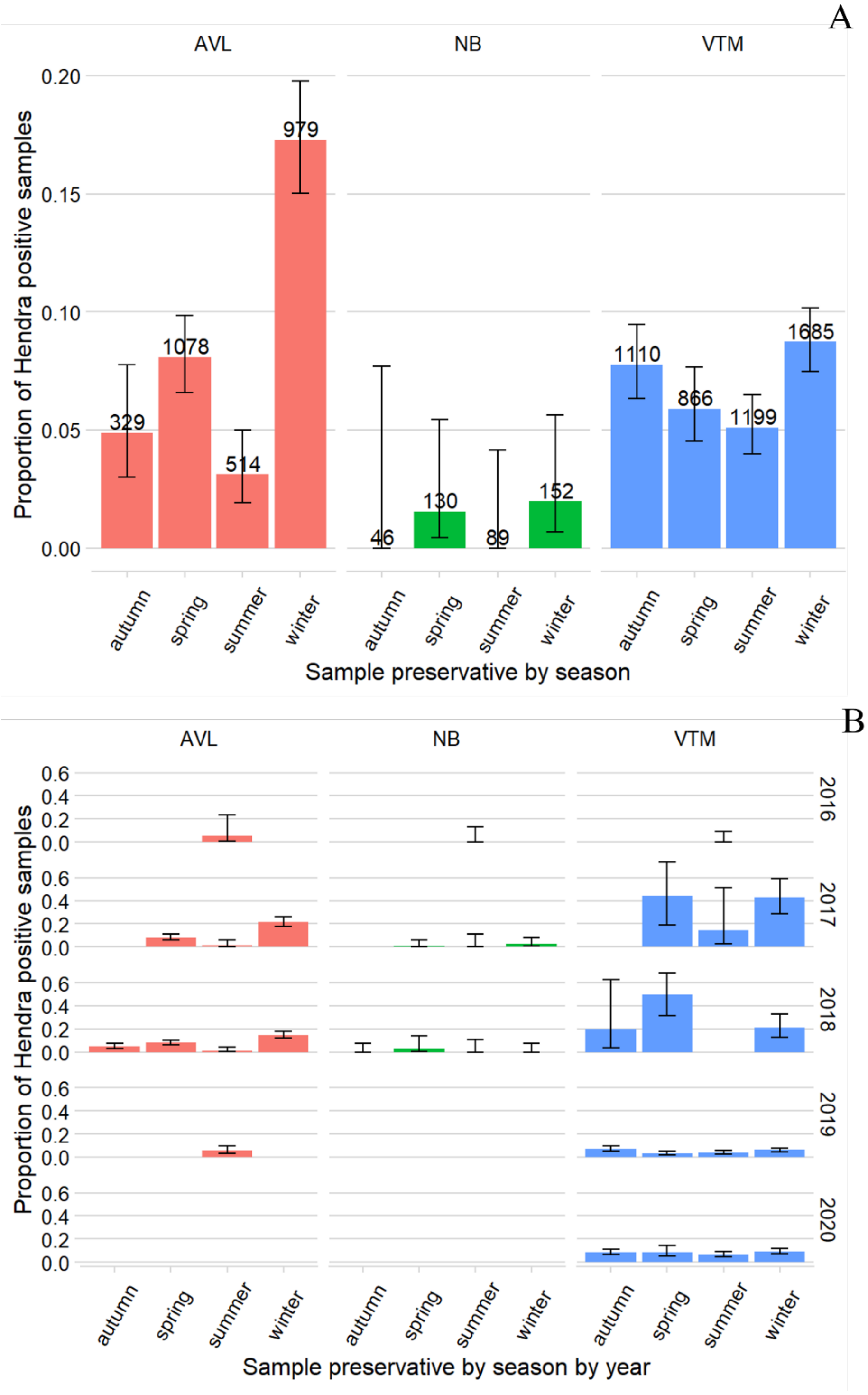


Figure S14: Effect of sample preservative on Hendra virus detection. A) shows samples overall, and B) samples split by year. Samples were aliquoted into an AVL lysis buffer (target 140ul of urine into 560ul buffer), a viral transport medium (VTM, between 200-1000µl of urine into 1000ul of VTM) or stored without a preservative (NB, up to 2000µl of urine) for Hendra virus testing.

Because of changes to the urine sample protocol – including a switch from AVL to VTM as the priority sample type – spatiotemporal patterns in shedding may be confounded by choice of sample buffer. To investigate this, we used occupancy modelling to estimate detection probabilities for each of the buffer types (AVL, VTM and no buffer, NB). Detection probabilities were estimated using the *unmarked* package, where repeat detections were represented by repeat tests on unique samples using different buffer types: 7,167 samples had only one buffer type tested, 415 had two buffer types tested, and 29 has all three buffer types tested. Estimated detection probabilities were: 0.825 for AVL, 0.555 for VTM, and 0.085 for no buffer. Detection from samples in VTM and no buffer were significantly lower than detection from samples in AVL (VTM: $p = 0.0413$, no buffer: $p=0.0000625$). Note - VTM is the most represented buffer in the database, with 4,822 samples, followed by AVL with 2,880 samples, then NB with 382. For analyses presented in the main text, permutations were done between samples of the same buffer type to account for bias introduced by preservation.

We acknowledge some limitations to interpreting proxies for viral load and prevalence from pooled samples. Pooled sampling is an efficient method for viral surveillance in Pteropodid bats because sampling of individual bats is time consuming, potentially hazardous and expensive, and detection rate of virus in individuals is very low (~4.2%, 95% CI 3.1–5.6%) (Edson et al. 2019). Pooled sampling across many individuals (with estimation of prevalence based on detection per pooled sample rather than per individual), was first described by Chua (2003) as a pathogen surveillance tool for *Pteropus* bats, and underpins all large scale sampling of Hendra virus to date (Field et al. 2011; Field et al. 2015; Edson et al. 2015; Burroughs et al. 2016). This approach may lead to overestimation of total prevalence if only one of many individuals contributing to the pool are infected (as described in Field et al. 2011; Giles et al. 2018). While this theoretically complicates interpretation of viral load of each individual contributing to the pooled sample, the impacts are expected to be low owing to its non-linear log scaling. In this study, we adopted an optimized pooled sampling approach to gain more realistic predictions of prevalence with minimum sampling bias and false negatives compared with previous under roost sampling methods (Giles et al. 2018). With these sampling amendments, screening of populations rather than individuals remains an effective approach to investigating Hendra virus infection dynamics in flying-fox populations (Field et al. 2011). To confirm patterns of shedding from pooled samples, and investigate bat-level drivers of shedding, future work could integrate data from individual bats (e.g., Hoegh et al. 2021). Alternatively, other methods of estimating prevalence, such as state-space or hidden Markov models, could be explored.

**Appendix S5: Isolation of Hendra virus relative to sample Ct**

Overall, the success of Hendra virus isolation from PCR-positive samples was low (11/239; 4.6%), but isolation from samples increased with higher viral load proxies. This relationship held true for samples from bats, horses and humans collectively (Figure S15A, Table S4), and for samples from bats only (Figure S15B, Table S4), from aggregated datasets (this study; Playford et al. 2010; Barr et al. 2015). When subdivided by dataset very few isolates were retained in the analysis (main dataset: 3/177, and historical dataset: 5/46, Table S4). Isolates of bat viruses are difficult to obtain: for Nipah virus – a closely related Henipavirus – isolation has only been successful on 17 occasions (Anderson et al. 2019); for bat coronaviruses, only five viral species have been successfully cultured (Ruiz-Aravena et al. 2022).

Note that isolations, but not Ct values, are described for samples in Playford et al. (2010) and Barr et al. (2015). Ct values relative to isolation from this study, Playford et al. (2010) and Barr et al. (2015) are given in table Table S5.

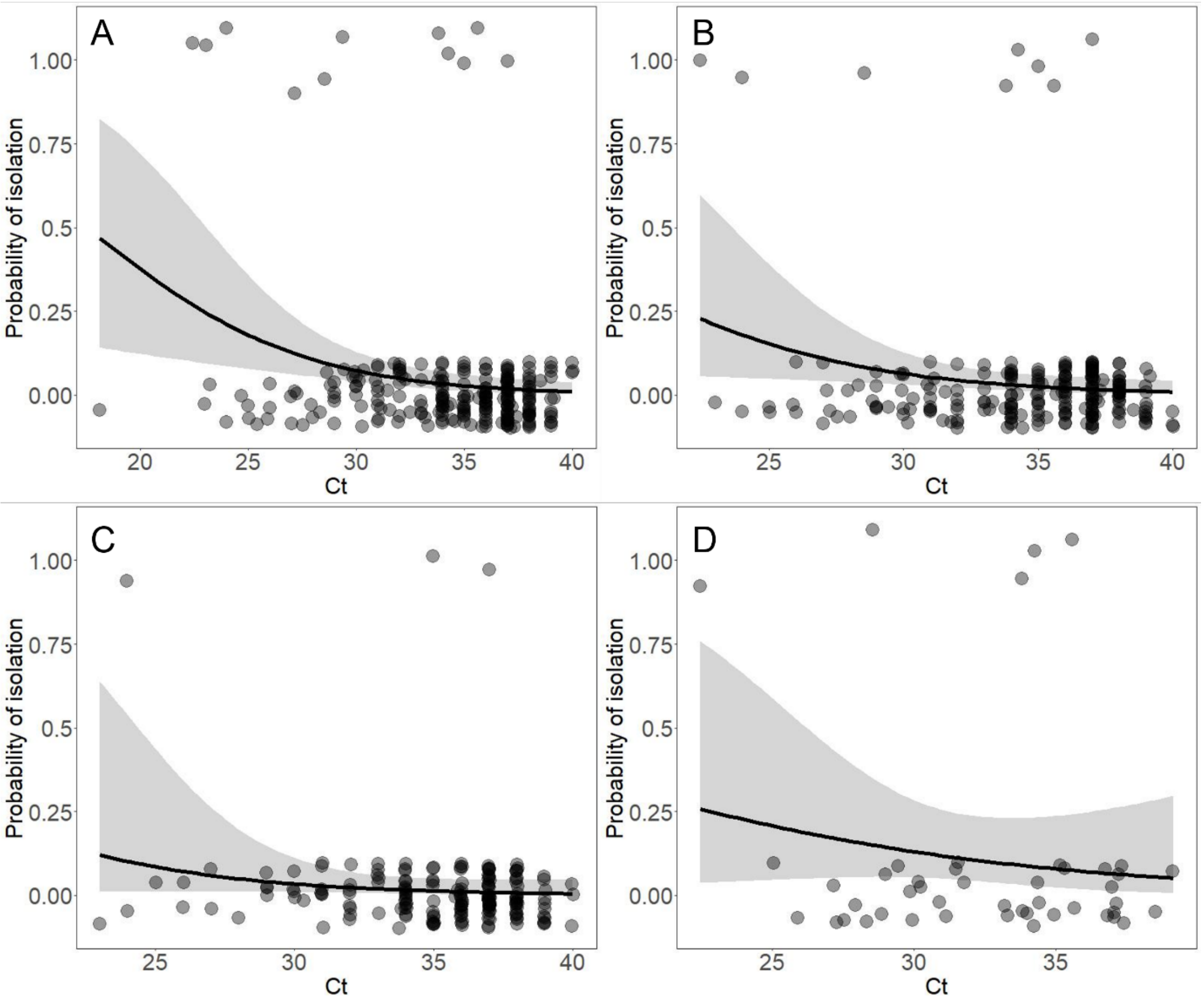


Figure S15: Logistic regression of isolation relative to Ct value of sample. (A) shows data from 239 Hendra positive samples taken from bats, horses and humans, with data aggregated from this study, Playford et al. (2010), and Barr et al. (2015), (B) from 223 Hendra positive samples taken from bats only from the aggregated datasets, (C) from 177 Hendra positive samples taken from bats only from this study, and (D) from 46 Hendra positive samples taken from bats only from Playford et al. (2010) and Barr et al. (2015). Plots show successful (top) and unsuccessful (bottom) isolation attempts. Isolation

Table S4: Logistic regression of isolation relative to Ct value of sample for separated datasets

| Dataset | Isolation | Coefficient (± SE) | p |
|---|---|---|---|
| Bats, horses and humans; aggregated datasets | 11/239; 4.6% | 18.4% ± 6.6% | 0.0014 |
| Bats only; aggregated datasets | 8/223; 3.6% | 17.6% ± 8.4% | 0.0164 |
| Bats only; this study | 3/177; 1.7% | 17.5% ±13.6% | 0.1299 |
| Bats only; Playford et al. 2010 & Barr et al. 2015 | 5/46, 10.9% | 10.5% ±12.4% | 0.3406 |

Table S5: Collated datasets on Hendra virus isolation. * indicates data from this study.

| **Isolation** | **Ct** | **Species** |
|---|---|---|
| 1 | 22.45 | flying-fox (individual) |
| 1 | 23.07 | human |
| 1 | 24 | *flying-fox (pooled) |
| 1 | 27.15 | horse |
| 1 | 28.55 | flying-fox (pooled) |
| 1 | 29.38 | horse |
| 1 | 33.8 | flying-fox (pooled) |
| 1 | 34.26 | flying-fox (pooled) |
| 1 | 35 | *flying-fox (pooled) |
| 1 | 35.6 | flying-fox (pooled) |
| 1 | 37 | *flying-fox (pooled) |
| 0 | 18.15 | human |
| 0 | 23 | *flying-fox (pooled) |
| 0 | 23.22 | human |
| 0 | 24 | *flying-fox (pooled) |
| 0 | 24.71 | human |
| 0 | 25 | *flying-fox (pooled) |
| 0 | 25.05 | flying-fox (individual) |
| 0 | 25.41 | human |
| 0 | 25.9 | flying-fox (pooled) |
| 0 | 26 | *flying-fox (pooled) |
| 0 | 26 | *flying-fox (pooled) |
| 0 | 27 | *flying-fox (pooled) |
| 0 | 27 | *flying-fox (pooled) |

| | | |
|---|---|---|
| 0 | 27.16 | flying-fox (pooled) |
| 0 | 27.26 | flying-fox (individual) |
| 0 | 27.53 | flying-fox (pooled) |
| 0 | 27.92 | flying-fox (pooled) |
| 0 | 28 | *flying-fox (pooled) |
| 0 | 28.32 | flying-fox (pooled) |
| 0 | 28.64 | horse |
| 0 | 28.84 | flying-fox (pooled) |
| 0 | 28.98 | flying-fox (pooled) |
| 0 | 29 | *flying-fox (pooled) |
| 0 | 29 | *flying-fox (pooled) |
| 0 | 29 | *flying-fox (pooled) |
| 0 | 29 | *flying-fox (pooled) |
| 0 | 29.44 | flying-fox (individual) |
| 0 | 29.87 | flying-fox (individual) |
| 0 | 29.95 | flying-fox (individual) |
| 0 | 30 | *flying-fox (pooled) |
| 0 | 30 | *flying-fox (pooled) |
| 0 | 30 | *flying-fox (pooled) |
| 0 | 30 | *flying-fox (pooled) |
| 0 | 30.15 | flying-fox (individual) |
| 0 | 30.24 | flying-fox (individual) |
| 0 | 30.29 | human |
| 0 | 30.34 | *flying-fox (individual) |
| 0 | 30.89 | flying-fox (pooled) |
| 0 | 31 | *flying-fox (pooled) |
| 0 | 31 | *flying-fox (pooled) |
| 0 | 31 | *flying-fox (pooled) |
| 0 | 31 | *flying-fox (pooled) |
| 0 | 31 | *flying-fox (pooled) |
| 0 | 31 | *flying-fox (pooled) |
| 0 | 31 | *flying-fox (pooled) |
| 0 | 31.03 | human |
| 0 | 31.13 | flying-fox (pooled) |
| 0 | 31.47 | flying-fox (individual) |
| 0 | 31.55 | flying-fox (pooled) |
| 0 | 31.78 | flying-fox (individual) |
| 0 | 31.95 | horse |
| 0 | 32 | *flying-fox (pooled) |
| 0 | 32 | *flying-fox (pooled) |
| 0 | 32 | *flying-fox (pooled) |
| 0 | 32 | *flying-fox (pooled) |
| 0 | 32 | *flying-fox (pooled) |
| 0 | 32 | *flying-fox (pooled) |

| | | |
|---|---|---|
| 0 | 32 | *flying-fox (pooled) |
| 0 | 32 | *flying-fox (pooled) |
| 0 | 32.12 | horse |
| 0 | 32.28 | human |
| 0 | 32.5 | horse |
| 0 | 33 | *flying-fox (pooled) |
| 0 | 33 | *flying-fox (pooled) |
| 0 | 33 | *flying-fox (pooled) |
| 0 | 33 | *flying-fox (pooled) |
| 0 | 33 | *flying-fox (pooled) |
| 0 | 33 | *flying-fox (pooled) |
| 0 | 33.19 | flying-fox (individual) |
| 0 | 33.31 | flying-fox (individual) |
| 0 | 33.78 | *flying-fox (individual) |
| 0 | 33.85 | flying-fox (pooled) |
| 0 | 33.87 | *flying-fox (individual) |
| 0 | 34 | *flying-fox (individual) |
| 0 | 34 | *flying-fox (pooled) |
| 0 | 34 | *flying-fox (pooled) |
| 0 | 34 | *flying-fox (pooled) |
| 0 | 34 | *flying-fox (pooled) |
| 0 | 34 | *flying-fox (pooled) |
| 0 | 34 | *flying-fox (pooled) |
| 0 | 34 | *flying-fox (pooled) |
| 0 | 34 | *flying-fox (pooled) |
| 0 | 34 | *flying-fox (pooled) |
| 0 | 34 | *flying-fox (pooled) |
| 0 | 34 | *flying-fox (pooled) |
| 0 | 34 | *flying-fox (pooled) |
| 0 | 34 | *flying-fox (pooled) |
| 0 | 34 | *flying-fox (pooled) |
| 0 | 34 | *flying-fox (pooled) |
| 0 | 34 | *flying-fox (pooled) |
| 0 | 34 | *flying-fox (pooled) |
| 0 | 34 | *flying-fox (pooled) |
| 0 | 34 | flying-fox (pooled) |
| 0 | 34.22 | flying-fox (pooled) |
| 0 | 34.38 | flying-fox (pooled) |
| 0 | 34.42 | flying-fox (pooled) |
| 0 | 34.56 | human |
| 0 | 34.94 | flying-fox (pooled) |
| 0 | 35 | *flying-fox (pooled) |
| 0 | 35 | *flying-fox (pooled) |
| 0 | 35 | *flying-fox (pooled) |

| | | |
|---|---|---|
| 0 | 35 | *flying-fox (pooled) |
| 0 | 35 | *flying-fox (pooled) |
| 0 | 35 | *flying-fox (pooled) |
| 0 | 35 | *flying-fox (pooled) |
| 0 | 35 | *flying-fox (pooled) |
| 0 | 35 | *flying-fox (pooled) |
| 0 | 35 | *flying-fox (pooled) |
| 0 | 35 | *flying-fox (pooled) |
| 0 | 35 | *flying-fox (pooled) |
| 0 | 35 | *flying-fox (pooled) |
| 0 | 35 | *flying-fox (pooled) |
| 0 | 35 | horse |
| 0 | 35.15 | flying-fox (pooled) |
| 0 | 35.32 | flying-fox (pooled) |
| 0 | 35.68 | flying-fox (pooled) |
| 0 | 35.81 | *flying-fox (individual) |
| 0 | 35.89 | *flying-fox (individual) |
| 0 | 36 | *flying-fox (pooled) |
| 0 | 36 | *flying-fox (pooled) |
| 0 | 36 | *flying-fox (pooled) |
| 0 | 36 | *flying-fox (pooled) |
| 0 | 36 | *flying-fox (pooled) |
| 0 | 36 | *flying-fox (pooled) |
| 0 | 36 | *flying-fox (pooled) |
| 0 | 36 | *flying-fox (pooled) |
| 0 | 36 | *flying-fox (pooled) |
| 0 | 36 | *flying-fox (pooled) |
| 0 | 36 | *flying-fox (pooled) |
| 0 | 36 | *flying-fox (pooled) |
| 0 | 36 | *flying-fox (pooled) |
| 0 | 36 | *flying-fox (pooled) |
| 0 | 36 | *flying-fox (pooled) |
| 0 | 36 | *flying-fox (pooled) |
| 0 | 36 | *flying-fox (pooled) |
| 0 | 36 | *flying-fox (pooled) |
| 0 | 36 | *flying-fox (pooled) |
| 0 | 36 | *flying-fox (pooled) |
| 0 | 36 | *flying-fox (pooled) |
| 0 | 36 | *flying-fox (pooled) |
| 0 | 36 | *flying-fox (pooled) |
| 0 | 36.76 | flying-fox (pooled) |
| 0 | 36.83 | flying-fox (pooled) |
| 0 | 36.98 | flying-fox (pooled) |
| 0 | 37 | *flying-fox (individual) |

| | | |
|---|---|---|
| 0 | 37 | *flying-fox (pooled) |
| 0 | 37 | *flying-fox (pooled) |
| 0 | 37 | *flying-fox (pooled) |
| 0 | 37 | *flying-fox (pooled) |
| 0 | 37 | *flying-fox (pooled) |
| 0 | 37 | *flying-fox (pooled) |
| 0 | 37 | *flying-fox (pooled) |
| 0 | 37 | *flying-fox (pooled) |
| 0 | 37 | *flying-fox (pooled) |
| 0 | 37 | *flying-fox (pooled) |
| 0 | 37 | *flying-fox (pooled) |
| 0 | 37 | *flying-fox (pooled) |
| 0 | 37 | *flying-fox (pooled) |
| 0 | 37 | *flying-fox (pooled) |
| 0 | 37 | *flying-fox (pooled) |
| 0 | 37 | *flying-fox (pooled) |
| 0 | 37 | *flying-fox (pooled) |
| 0 | 37 | *flying-fox (pooled) |
| 0 | 37 | *flying-fox (pooled) |
| 0 | 37 | *flying-fox (pooled) |
| 0 | 37 | *flying-fox (pooled) |
| 0 | 37 | *flying-fox (pooled) |
| 0 | 37 | *flying-fox (pooled) |
| 0 | 37 | *flying-fox (pooled) |
| 0 | 37 | *flying-fox (pooled) |
| 0 | 37 | *flying-fox (pooled) |
| 0 | 37 | *flying-fox (pooled) |
| 0 | 37 | *flying-fox (pooled) |
| 0 | 37 | *flying-fox (pooled) |
| 0 | 37 | *flying-fox (pooled) |
| 0 | 37 | *flying-fox (pooled) |
| 0 | 37 | *flying-fox (pooled) |
| 0 | 37 | *flying-fox (pooled) |
| 0 | 37 | *flying-fox (pooled) |
| 0 | 37 | *flying-fox (pooled) |
| 0 | 37 | *flying-fox (pooled) |
| 0 | 37.07 | flying-fox (pooled) |
| 0 | 37.08 | flying-fox (pooled) |
| 0 | 37.15 | flying-fox (pooled) |
| 0 | 37.23 | flying-fox (pooled) |
| 0 | 37.32 | flying-fox (pooled) |
| 0 | 37.41 | flying-fox (pooled) |
| 0 | 38 | *flying-fox (individual) |
| 0 | 38 | *flying-fox (individual) |

| | | |
|---|---|---|
| 0 | 38 | *flying-fox (individual) |
| 0 | 38 | *flying-fox (individual) |
| 0 | 38 | *flying-fox (individual) |
| 0 | 38 | *flying-fox (individual) |
| 0 | 38 | *flying-fox (individual) |
| 0 | 38 | *flying-fox (pooled) |
| 0 | 38 | *flying-fox (pooled) |
| 0 | 38 | *flying-fox (pooled) |
| 0 | 38 | *flying-fox (pooled) |
| 0 | 38 | *flying-fox (pooled) |
| 0 | 38 | *flying-fox (pooled) |
| 0 | 38 | *flying-fox (pooled) |
| 0 | 38 | *flying-fox (pooled) |
| 0 | 38 | *flying-fox (pooled) |
| 0 | 38 | *flying-fox (pooled) |
| 0 | 38 | *flying-fox (pooled) |
| 0 | 38 | *flying-fox (pooled) |
| 0 | 38 | *flying-fox (pooled) |
| 0 | 38 | *flying-fox (pooled) |
| 0 | 38 | *flying-fox (pooled) |
| 0 | 38 | *flying-fox (pooled) |
| 0 | 38 | *flying-fox (pooled) |
| 0 | 38 | *flying-fox (pooled) |
| 0 | 38.53 | flying-fox (pooled) |
| 0 | 39 | *flying-fox (individual) |
| 0 | 39 | *flying-fox (pooled) |
| 0 | 39 | *flying-fox (pooled) |
| 0 | 39 | *flying-fox (pooled) |
| 0 | 39 | *flying-fox (pooled) |
| 0 | 39 | *flying-fox (pooled) |
| 0 | 39 | *flying-fox (pooled) |
| 0 | 39 | *flying-fox (pooled) |
| 0 | 39 | *flying-fox (pooled) |
| 0 | 39 | *flying-fox (pooled) |
| 0 | 39 | *flying-fox (pooled) |
| 0 | 39.15 | flying-fox (pooled) |
| 0 | 40 | *flying-fox (pooled) |
| 0 | 40 | *flying-fox (pooled) |
| 0 | 40 | *flying-fox (pooled) |
